# Suppression of HSF1 activity by wildtype p53 creates the driving force for p53 loss-of-heterozygosity, enabling mutant p53 stabilization and invasion

**DOI:** 10.1101/2020.04.23.057034

**Authors:** Özge Cicek Sener, Adrian Stender, Luisa Klemke, Nadine Stark, Tamara Isermann, Jinyu Li, Ute M. Moll, Ramona Schulz-Heddergott

**Affiliations:** Institute of Molecular Oncology, University Medical Center Göttingen, 37077 Göttingen, Germany; Department of Pathology, Stony Brook University, Stony Brook NY 11794, USA

**Keywords:** mutp53, HSF1, Hsp90, Hsp70, CDK4, MLK3, MAPK, AOM/DSS, colorectal cancer, organoids, Idasanutlin

## Abstract

A prerequisite for gain-of-function (GOF) p53 missense mutants (mutp53) is protein stabilization. Moreover, a prerequisite for mutp53 stabilization is loss of the remaining wildtype (WT) p53 allele (loss-of-heterozygosity, p53LOH) in mutp53/+ tumors. Thus, GOF, mutp53 stabilization and p53LOH are strictly linked. However, the driving force for p53LOH is unknown. Typically, heterozygous tumors are an instable transition state. Here we identify the repressive WTp53-HSF1 axis as the driver of p53LOH.

We find that the WTp53 allele in AOM/DSS-induced colorectal tumors (CRC) of p53^R248Q/+^ mice retains its haploid transcriptional activity. Notably, WTp53 represses heat-shock factor 1 (HSF1) activity, the master transcription factor of the proteotoxic stress defense response (HSR) that is ubiquitously and constitutively activated in cancer tissues. HSR is critical for stabilizing oncogenic proteins including mutp53. WTp53-retaining murine CRC tumors and tumor-derived organoids and human CRC cells all suppress the tumor-promoting HSF1 transcriptional program.

Mechanistically, the retained WTp53 allele activates CDKN1A/p21, leading to cell cycle inhibition and suppression of the E2F target gene MLK3. MLK3 links cell cycle to the MAPK stress pathway to activate the HSR response. We show that in p53^R248Q/+^ tumors WTp53 activation by constitutive stress (emanating from proliferative/metabolic stresses and genomic instability) represses MLK3, consequently inactivating the MAPK-HSF1 response necessary to ensure tumor survival. This creates strong selection pressure for p53LOH which eliminates the repressive WTp53-HSF1 axis and unleashes the tumor-promoting HSF1 functions, inducing mutp53 stabilization and enabling invasion.

**HIGHLIGHTS:**

- heterozygous p53^R248Q/+^ tumors retain p53 transcriptional activity in a mouse model of colorectal cancer (CRC)
- wildtype p53 actively represses the tumor-promoting HSF1-regulated chaperone system and proteotoxic stress response
- the repressive WTp53 – HSF1 axis creates a selective pressure for WTp53 loss-of-heterozygosity in CRC tumors
- p53 loss-of-heterozygosity enables stabilization of the gain-of-function p53^R248Q^ mutant protein which in turn enables CRC invasion

## Introduction

Colorectal cancer (CRC) is due to several driver mutations and the third leading cause of cancer deaths worldwide. TP53 mutations enable the critical transition from late adenoma to invasive carcinoma^1, 2^. Next to APC, TP53 mutations are the second most common alteration in sporadic CRC, affecting > 60% of cases^3-10^. The vast majority of TP53 alterations are missense mutations (mutp53) with hotspot codons R175, G245, R248, R273 and R282^11-13^. In addition to loss-of-WTp53 function (LOF), some, especially hotspot, mutp53 alleles gain broad tumorigenic gain-of-function (GOF) and actively promote aggressive cancer progression *in vivo*^9, 14-20^. Some GOF mutants acquire allele-specific functions, not necessarily shared by other mutants^3, 9, 21-26^. The GOF *TP53*^*R248Q*^ allele is one of the most common across cancer types^10^.

A prerequisite for GOF is the tumor-specific stabilization of mutp53 proteins by the HSP90/HSP70/HSP40 chaperone systems^27-31^, providing protection from degradation by E3-ubiquitin ligases Mdm2 and CHIP^32, 33^. HSF1, the master transcription factor of the inducible heat-shock stress response (HSR), governs stress-induced chaperones including HSP90, HSP70 and HSP40 and is the major proteotoxic defense in tumors, preventing aberrant oncoproteins from aggregation^34-36^. Moreover, HSF1 induces chaperone-independent tumor-promoting genes, together imparting on HSF1 a key co-oncogenic role in tumorigenesis^37-39^. Notably, since cancer cells experience cumulative stress during tumorigenesis, HSF1 is increasingly activated^40^.

p53LOH is a critical prerequisite for mutp53 stabilization in tumors. Heterozygous tumors rarely if ever stabilize p53 *in vivo*^9, 19, 41, 42^. Importantly, the majority of human mutp53 tumors have undergone p53LOH^43-47^. Moreover, p53LOH has watershed significance in promoting tumor progression. Recent mouse studies clearly identify p53LOH as strong tumor promoting force^41, 48, 49^. Our previous studies comparing sarcomas and breast cancer identified that heterozygous mutp53 tumors require a second hit for mutp53 stabilization, i.e. loss of the remaining WTp53 allele^41^. However, the driving force behind p53LOH remained elusive.

Given how important mutp53 GOF activities are in tumor biology, it is imperative to understand the mechanism that drives stabilization of GOF mutants. The dependency of mutp53 stabilization on p53LOH appears somehow regulated by the remaining WTp53, but its mechanism is unknown. Using a genetically controlled p53LOH system in a CRC model, we show their causal relationship. We identify that the remaining WTp53 allele in p53^R248Q/+^ tumors represses the HSF1 chaperone axis, thereby preventing mutp53^R248Q^ protein stabilization, GOF and invasion. This creates a strong driving force for p53LOH. In sum, a single pivotal genetic event, p53LOH, simultaneously provides three major evolutionary forces to drive cancer, *i*) loss of residual WTp53 suppressor activity including the repressive WTp53-HSF1 axis, *ii)* tumor-promoting HSF1 upregulation, and *iii*) mutp53 protein stabilization which liberates GOF activities. This provides an explanation for the longstanding puzzle why p53LOH strictly correlates with mutp53 stabilization and higher tumor aggressiveness.

## RESULTS

### p53LOH is a prerequisite for mutp53 stabilization and invasion in colorectal cance

Stabilization of missense mutant p53 (mutp53) proteins specifically in tumor but not normal cells is a key feature and prerequisite of GOF^16, 42^. Since p53LOH is a critical prerequisite for mutp53 stabilization in sarcomas and breast cancer^41^, we examined mutp53 stabilization before and after p53LOH in the colorectal AOM/DSS model^9^. Briefly, we combined the humanized GOF *TP53*^R248Q^ allele (short ‘p53^Q^’) with either p53 wildtype (WT, ‘+’) or knock-out (‘-’) alleles and determined the p53LOH effect on mutp53 levels (Figures S1A-B). Indeed, massive mutp53 stabilization was detected in 100% of p53^Q/-^ tumors, whereas 100% of p53^Q/+^ tumors failed to undergo stabilization (Figure S1B). Notably, p53LOH increases tumor numbers in the p53^-/-^ and even more so in the p53^Q/-^ setting (Figure S1C). Notably, 100% of tumors retaining one WTp53 allele (p53^Q/+^ and p53^-/+^) remain noninvasive (Figure S1D). Conversely, loss of the remaining WTp53 allele (p53^Q/-^ or p53^-/-^) enables invasion (Figures S1D-E). Thus, p53LOH is the critical determinant for CRC invasion.

To independently validate that p53LOH enables mutp53 stabilization and CRC invasion, we used a second inducible model that combines the constitutive p53^Q^ allele with a floxed WTp53 (p53^fl^) allele. p53^Q/fl^ mice were crossed to *villinCreER*^*T2*^ mice to generate Tamoxifen (TAM)-inducible p53LOH restricted to intestinal epithelial cells (p53^Q/Δ^), plus non-LOH controls (p53^Q/+^) (Figure 1A). Importantly, TAM-mediated p53LOH was induced uniformly at a defined tumor burden verified by colonoscopy (Figure 1B)^50^. Controls were (*i*) p53^Q/fl^ oil-treated mice, and (*ii*) p53^Q/+^ TAM-treated mice to exclude nonspecific TAM effects. At 6-8 wks post TAM, LOH tumors showed a trend towards increased tumor numbers and sizes compared to both ‘no LOH’ control groups (Figures 1C-D). When analyzed earlier at 3-5 wks post TAM, tumor burden had not yet increased, indicating that LOH’s effect on promoting proliferation requires time and is incremental (Figure 1E).

**Figure 1.**
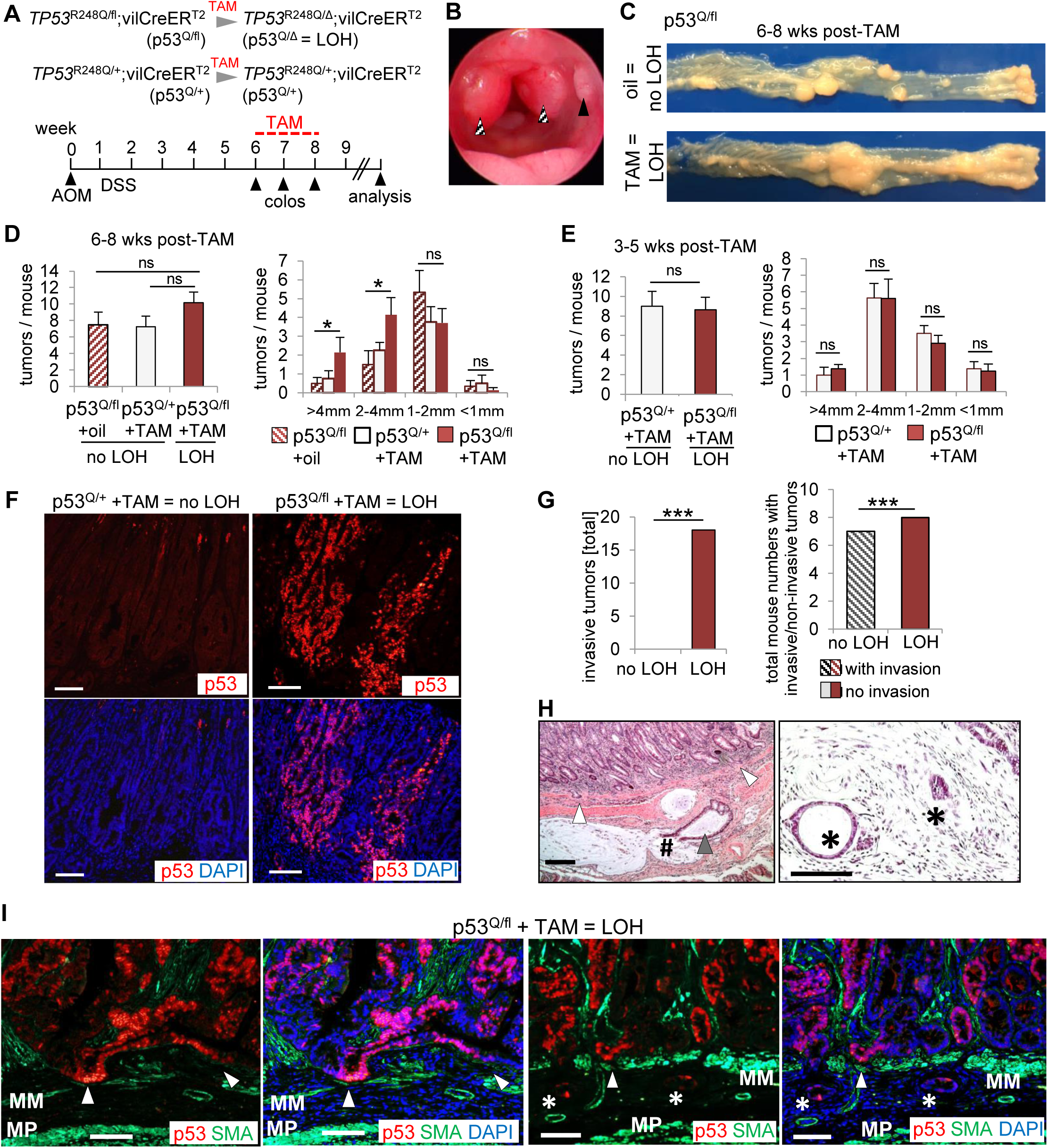
p53 loss-of-heterozygosity (p53LOH) is a prerequisite for mutp53 protein stabilization and enables invasion in colorectal cancer. (A) p53LOH induction scheme in intestinal epithelial cells. The constitutive GOF TP53^R248Q^ allele (p53^Q^) is paired with the conditional wildtype Trp53 allele (p53^fl^) harboring loxP sites in Introns 2 and 10 for Tamoxifen (TAM)-induced Cre recombinase-mediated deletion. A constitutive WTp53 allele (‘p53^+^’) serves as control. Colorectal tumors were initiated by a bolus of AOM/DSS at age 10 wks. Tumor burden was visualized by weekly colonoscopy (colos). At a defined tumor burden (when heterozygous mice had at least 2-3 S2 tumors and at least one S3 tumor), TAM or oil treatment was administered. Oil-treated p53^Q/fl^;vilCreER^T2^ and TAM-treated p53^Q/+^;vilCreER^T2^ control mice carry heterozygous tumors, whereas TAM-treated TP53^R248Q/fl^;vilCreER^T2^ (p53^Q/Δ^) mice carry p53LOH tumors. TAM-treated TP53^R248Q/+^;vilCreER^T2^ mice served as additional control to exclude nonspecific TAM effects. Mice were analyzed 2-8 wks after LOH induction. (B) Representative colonoscopy image of an untreated TP53^R248Q/fl^;vilCreER^T2^ mouse at 8 wks post AOM. Tumors with scores S2 (solid arrow) and S3 (dashed arrows)^50^. (C) Representative macroscopic view of the entire dissected colons of p53^Q/fl^ mice treated as indicated, 6 wks after inducing p53LOH. Left end, ileocecal valve; right end anus. (D) Number of colonic tumors per mouse (left) and tumor size distribution (right) of the indicated genotypes analyzed at 6-8 wks post TAM or oil treatment. Note that both ‘no LOH’ groups had the same tumor burden. p53^Q/fl^ + oil group with n=6, p53^Q/+^ + TAM group with n=5, p53^Q/fl^ + TAM group with n=8. Mean ± SEM, Student’s t-test. p*=0.05; ns, not significant. (E) Number of colonic tumors per mouse (left) and tumor size distribution (right) of the indicated genotypes analyzed earlier than in (D) at 3-5 wks post TAM. p53^Q/+^ + TAM group with n=8, p53^Q/fl^ + TAM group with n=13. Mean ± SEM, Student’s t-test. ns, not significant. (F) Representative immunofluorescence of p53 for TAM-treated p53^Q/+^ (‘no LOH’) and TAM-treated p53^Q/fl^ mice (‘LOH’) at endpoint 6 wks post-TAM. Scale bars, 100 μm. (G) Total number of invasive tumors (left) and numbers of mice with non-invasive and invasive tumors and (right) of oil-treated p53^Q/fl^ mice and TAM-treated p53^Q/+^ mice (combined as ‘no LOH’ group) versus TAM-treated p53^Q/fl^ mice (‘LOH’ group) analyzed at 6-8 wks post oil/TAM. (left) ‘no LOH’, n = 27 tumors from 7 mice analyzed and ‘LOH’, n = 49 tumors from 8 mice analyzed. (right) ‘no LOH’ n = 7 mice and ‘LOH’ n = 8 mice. Fisher’s exact test. Bars, mean ± SEM. p***≤ 0.001. (H) Representative histopathology of two LOH p53^Q/fl^ tumors 8 wks after TAM treatment. LOH induction showing (left) extensive invasion deep within the muscularis propria of the bowel wall (black arrow) and the muscularis mucosae (white arrows). The deeply invasive malignant gland (black arrow) is ruptured, spilling its content into mucous lakes with single tumor cells floating in it (#). Right, invading glands in the submucosa (*). H&E, scale bars 100 µm. (I) Representative immunofluorescence of two invasive tumors stabilized for mutp53 from TAM-treated p53^Q/fl^ mice (‘LOH’ group) for p53 (red), α-SMA (smooth muscle marker, green) and DAPI (blue) at endpoint 6 wks. Scale bars, 100 μm. MM, muscularis mucosae, MP, muscularis propia. White arrows, tumor cells invading the MM. Asterix, small invasive cell clumps invading the submucosa. Note that tips of invading cells are predominantly positive for mutp53.

Mutp53 stabilization is a critical prerequisite for mutp53 GOF^9^. Indeed, all p53LOH mice exhibited stabilized mutp53, in sharp contrast to mice with a retained WTp53 allele (Figure 1F). The WTp53 allele is a major barrier to tumor invasion as reported by us and others^7, 9, 42^. In agreement, ‘no LOH’ mice (oil-treated p53^Q/fl^ and TAM-treated p53^Q/+^ mice) never developed invasive tumors (Figure 1G). In stark contrast, induced p53LOH caused a dramatic increase in invasive tumors in the cohort from 0/27 tumors to 18/49 tumors (Figures 1G-I) and all mice harbored at least one invasive tumor. Notably, mutp53 stabilization is particularly prominent at the invasive front (Figure 1I). Moreover, in the constitutive p53LOH model, tumors lacking WTp53 confirmed the dramatic increase in invasion (Figures S1D-E). In sum, while LOH only has an incremental effect on tumor proliferation, p53LOH is a dramatic gate-opener unleashing GOF by mediating mutp53 stabilization, which in turn enables invasion.

### The WTp53 allele in heterozygous colorectal tumors retains its activity and suppresses the HSF1 transcriptional program

While mutp53 stabilization after p53LOH is dramatic, the mechanism of tumor-specific mutp53 accumulation triggered by p53LOH is incompletely understood. In agreement with other studies^16, 51^, loss of Mdm2 induction by the WT allele might play some role (Figure 2A, compare Mdm2 mRNA in p53^Q/+^ *vs* p53^Q/-^ tumors). However, an additional mechanism likely exists to ensure such massive stabilization after p53LOH.

**Figure 2.**
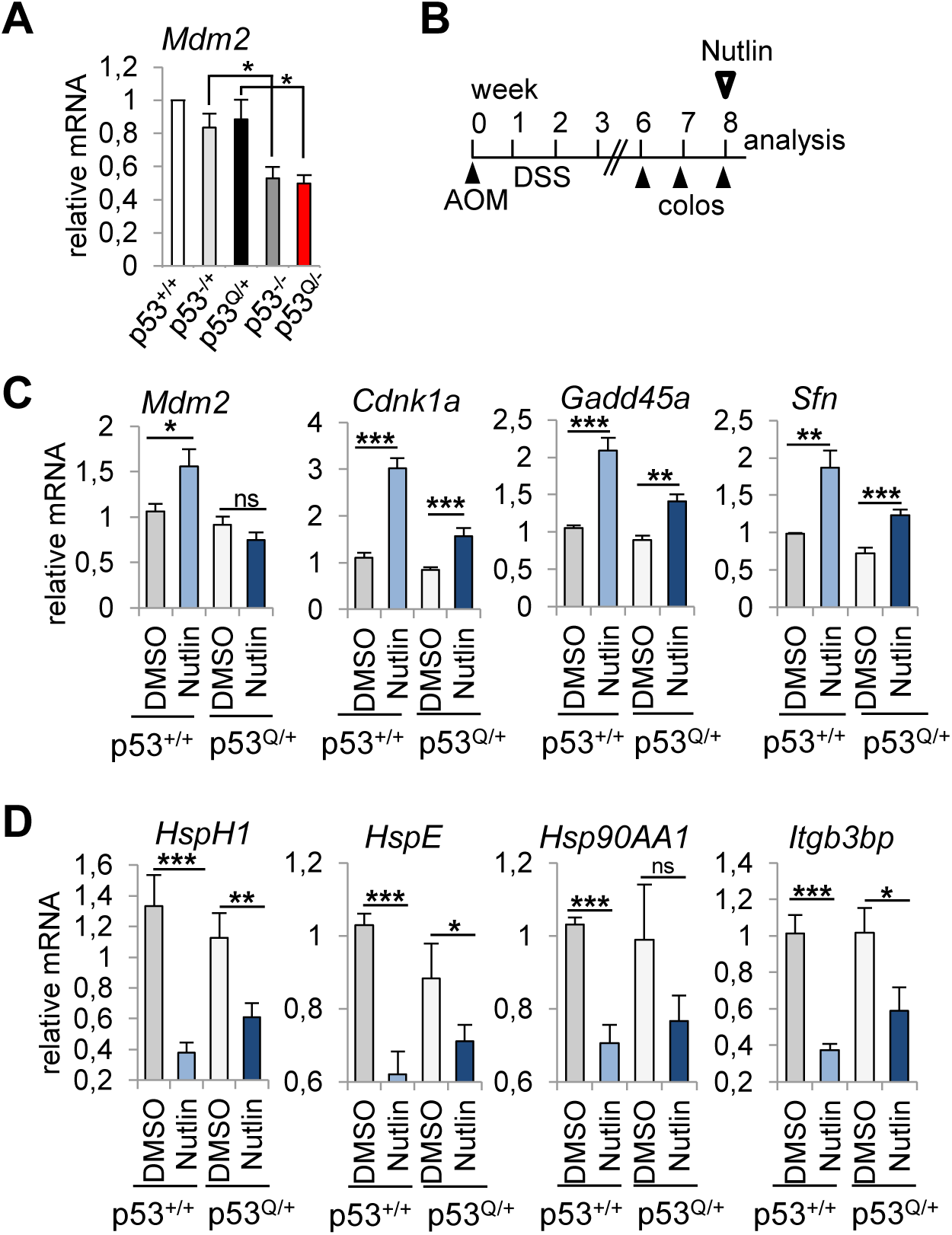
The WTp53 allele in heterozygous colorectal tumors retains its activity and represses HSF1 target gene expression in vivo. (A) Mdm2 mRNA levels of untreated CRC tumors from the indicated genotypes. Single colonic tumors from the indicated genotypes were pooled (≥ 5 tumors per group). qRT-PCR normalized to 36B4 mRNA. Mean ± SEM of 3 technical replicates, each in triplicates. Student’s t-test. (B) Scheme of Nutlin treatment in the AOM/DSS colorectal tumor model. After AOM/DSS induction, tumor growth was quantitated by serial colonoscopy (‘colos’). Mice with the defined tumor burden of at least 2-3 S2 tumors and at least one S3 tumor were orally treated with vehicle or 150mg/kg Nutlin for 3 days. Tumors were analyzed 8 hrs after the last treatment. (C, D) mRNA levels of WTp53 target genes (C) or HSF1 target genes (D) of colonic p53^Q/+^ and p53^+/+^ tumors of DMSO- and Nutlin-treated mice. Single colonic tumors from the indicated genotypes were pooled (≥ 5 tumors per group). qRT-PCR normalized to 36B4 mRNA. Mean ± SEM of 3 technical replicates, each in triplicates. Student’s t-test.

A major pathway for tumor-specific mutp53 stabilization is the intrinsic tumor stress-induced HSF1-governed chaperone system^14, 33, 35, 52, 53^. In cancer cells the constitutively (phospho-) activated master transcription factor HSF1 orchestrates the major proteotoxic defense. Thus, we asked whether in heterozygous tumors the remaining WTp53 suppresses global HSF1 activity or distinct chaperone targets. This hypothesis assumes that despite the presence of a GOF allele (Q in this case), the remaining WTp53 allele at least partially retains its transcriptional activity. Thus, we treated tumor-bearing p53^Q/+^ mice with Nutlin, a highly specific non-genotoxic p53 activator inhibiting its E3 ligase MDM2, to mimic the general activation state of WTp53 in tumors constitutively stressed by aberrant growth and metabolic stress, hypoxia and genomic instability (Figure 2B).

Indeed, in p53^Q/+^ tumors Nutlin induced allele-dose dependently (haploid) WTp53 target gene expression (e.g. Cdnk1a, Gadd45a and Sfn) (Figure 2C). Interestingly, however, Mdm2 expression after Nutlin only increased in p53^+/+^ but not in p53^Q/+^ tumors (Figure 2C), indicating that p53-regulated Mdm2 levels cannot account for the missing mutp53 stabilization in heterozygous tumors. Why Mdm2 failed to increase remains unclear. We conclude that, surprisingly, the GOF mutp53^R248Q^ allele fails to exert a dominant-negative effect over the remaining WTp53 allele as predicted by many, mainly *in vitro*, studies^54-57^. Importantly, this residual WTp53 activity is sufficient to suppress canonical HSF1 target genes in p53^Q/+^ tumors (Figure 2D). As expected from the double allelic dose, p53^+/+^ tumors showed a stronger HSF1 target gene suppression after Nutlin (Figure 2D).

We next tested whether simple loss of the WTp53 allele is able to activate HSF1 without Nutlin. While p53-/- vs. p53+/+ AOM/DSS mice have accelerated tumor growth (larger tumor numbers and sizes, Figures S2A-D) due to reduced cell cycle inhibitory/pro-apoptotic p53 target gene expression (Figures S2E-F), only some HSF1 target genes increased (Figure S2G). Conversely, stress-activated WTp53, mimicked by Nutlin, suppresses HSF1 activity to prevent chaperone-mediated mutp53 stabilization (Figure 2D).

In sum, in a stressed tumor milieu activated WTp53 in heterozygous mutp53/+ tumors creates the driving force for p53LOH. p53LOH eliminates the repressive WTp53-HSF1 axis and enables activation of the broad co-oncogenic HSF1 functions, which causes mutp53 protein stabilization that in turn enables tumor growth but foremost invasion.

### Activated WTp53 represses HSF1 activity in human colorectal cancer cells

Since mutp53 stabilization specifically arises in the malignant epithelial compartment, we analyzed the mechanism of p53-mediated HSF1 suppression in human CRC cell lines harboring WTp53. We resorted to homozygous WTp53 lines because heterozygous human CRC lines are not readily available. Importantly, measuring the global HSF1-mediated HSR response by heat-shock response element (HSE) luciferase assay confirmed HSF1 suppression upon WTp53 activation by Nutlin (Figure 3A). Moreover, Nutlin-induced HSF1 suppression was rescued by shp53-mediated depletion, confirming that the Nutlin-induced effect is p53-specific (Figure 3B).

**Figure 3.**
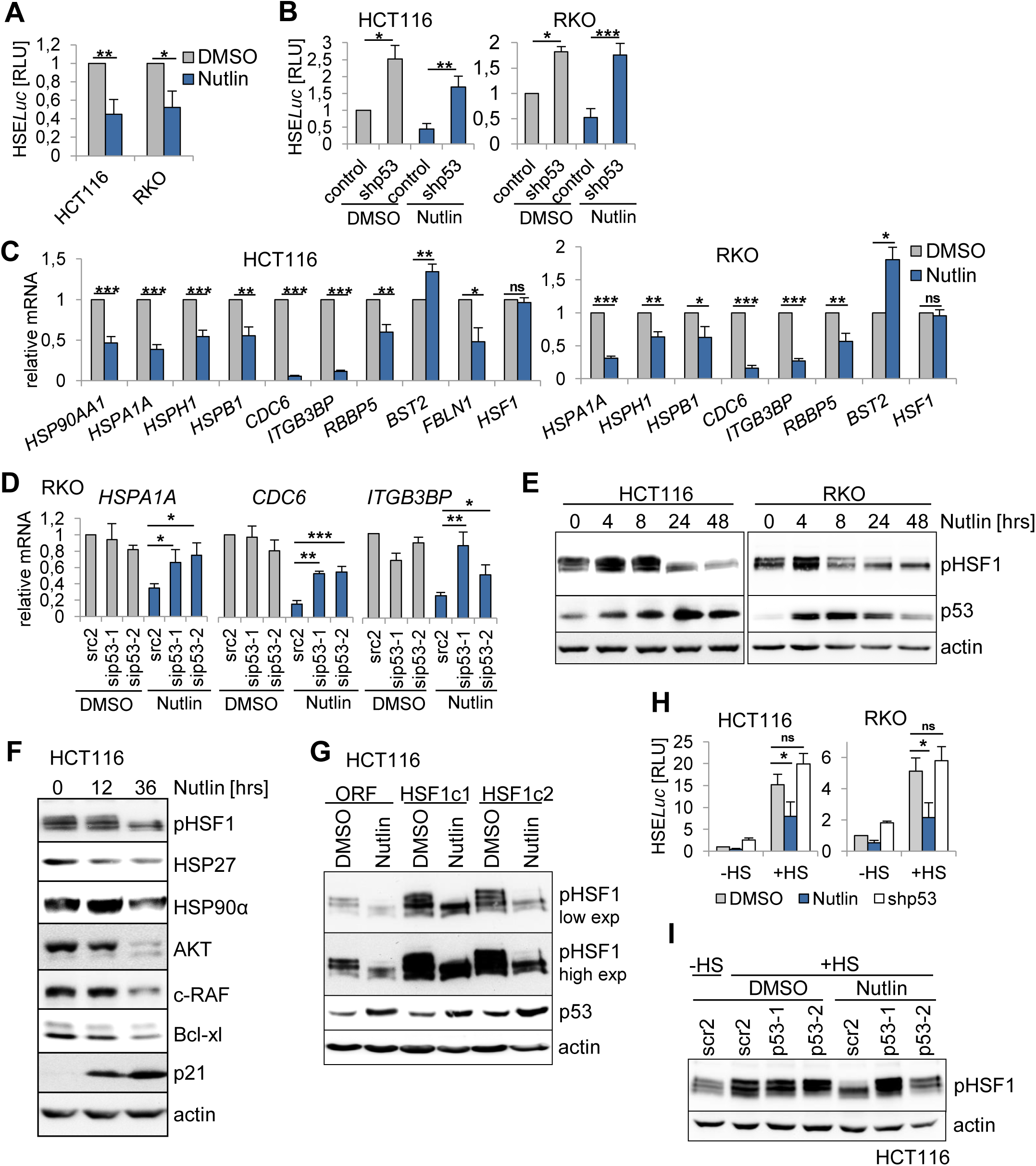
Activated WTp53 represses HSF1 activity in human colorectal cancer cells. (A) Luciferase reporter assay for heat-shock response elements (HSE). HCT116 and RKO cells were co-transfected with HSE-*Luc* and *Renilla* plasmids (pRL-TK). 48 hrs later cells were treated with DMSO or 10 µM Nutlin for 24 hrs. *Firefly* expression was normalized to *Renilla* expression and relative light units (RLU) were calculated. Mean ± SEM of 3 independent experiments, each in triplicates. Student’s t test. (B) HSE Luciferase assay as in (A) upon depletion of WTp53 by shRNA. Control, scramble shRNA. 48 hrs post transfection, cells were treated +/- Nutlin (10 µM) for 24 hrs. HSF1 binding to its HSE-promoter was measured as in (A). Mean ± SEM of 3 independent experiments, each in triplicates. Student’s t test. (C) Chaperone-dependent and -independent HSF1 target gene expression in HCT116 and RKO cells treated with DMSO or 10 µM Nutlin for 24 hrs. qRT-PCR for the indicated mRNAs, each normalized to 36B4 mRNA. Relative values are given in [ratio (2^-ddCT^)]. Mean ±SEM of 2 independent experiments, each repeated twice in triplicates. Student’s t-test, p*=0.05, p**=0.01, p***=0.001; ns, not significant. (D) HSF1 target gene expression in RKO cells upon depletion of WTp53. 48 hrs post transfection with two sip53 RNAs or scrambled control siRNA (scr2), RKO cells were treated +/- Nutlin (10 µM) for 24 hrs. qRT-PCR for the indicated mRNAs as in (C). Mean ±SEM of 2 independent experiments, each repeated twice in triplicates. Student’s t-test, p*=0.05, p**=0.01, p***=0.001. (E) Activated WTp53 suppresses pSer326-HSF1, the key marker of HSF1 activity. HCT116 and RKO cells were treated +/- Nutlin (10 µM) for the indicated times. Immunoblot analysis. p53 accumulation indicates p53 activation. Actin, loading control. (F) Repression of HSF1 target genes (Hsp90α and Hsp27) and destabilization of the Hsp90α client proteins AKT, c-Raf and Bcl-xl after p53 activation. HCT116 cells were treated with DMSO or 10 µM Nutlin for the indicated times. Immunoblot analysis. Actin, loading control. (G) Stably HSF1-overexpressing HCT116 subclones (HSF1c1 and HSF1c2) or empty vector control cells (ORF) were treated with DMSO or Nutlin for 24 hrs. Representative immunoblot analysis. pSer326-HSF1 shown with short and long exposure times. Actin, loading control. (H) Nutlin represses HSF1 activity in heat-shocked cells, rescued by p53 knockdown. HSE luciferase assay. HCT116 and RKO cells were transfected with HSE-*Luc* and *Renilla* plasmids and shp53 as in (B). 48 hrs post transfection, cells were treated with DMSO or 10 µM Nutlin for 24 hrs. During the final 2 hrs, HCT116 cells were heat-shocked for 1 hr at 42°C followed by recovery for 1 hr. HSF1 binding to HSE-*Luc* reporter was measured as in (A). Mean ± SEM of 3 independent experiments, each in triplicates. Student’s t-test, p*=0.05; ns, not significant. (I) The heat-shock response is markedly attenuated by Nutlin, while p53 depletion rescues Nutlin-induced HSF1 inactivation. HCT116 cells were transfected with different sip53 RNAs or scrambled (scr2). 48 hrs post transfection, cells were treated with DMSO or 10 µM Nutlin for 24 hrs. During the final 2 hrs, HCT116 cells were heat-shocked for 1 hr at 42°C followed by recovery for 1 hr as in (H). Immunoblot analysis for pSer326-HSF1. Actin, loading control.

HSF1 not only orchestrates the cellular chaperone system. In cancer cells HSF1 also broadly upregulates a large palette of tumor-promoting genes involved in cell cycle, DNA repair, metabolism, adhesion and protein translation^34^. Thus, we analysed randomly selected HSF1 targets representing different functions (Figure 3C). Notably, upon p53 activation by Nutlin we observed repression of classic HSF1 targets including *HSP90AA1, HSPA1A, HSPH1* and *HSPB1*, validating the mouse model (Figure 3C). Moreover, Nutlin also suppressed the tumor-promoting HSF1 targets *CDC6, ITGB3BP, RBBP5, BST2* and *FBLN1* (Figure 3C). Importantly, p53 depletion by siRNAs rescued their repression, confirming that p53 specifically regulates HSF1 activity (Figures 3D, S3A). The critical phosphorylation site for HSF1 activation is residue Ser326 which serves as functional hallmark of the tumor-promoting HSR response^37, 58^. Concomitantly to HSF1 target gene repression, p53 activation profoundly reduced pSer326-HSF1 levels in HCT116, RKO (Figure 3E), LS513 and LS174T cells (Figure S3B). Again, p53 depletion by siRNAs (Figure S3C) or p53 deletion in isogenic HCT116 cells (Figure S3D) abolished pSer326-HSF1 dephosphorylation. Conversely, mutp53-haboring CRC cells failed to repress HSF1 after Nutlin (Figure S3E). Of note, total HSF1 protein remained unchanged, excluding that HSF1 dephosphorylation/ inactivation is simply a consequence of reduced total HSF1 levels (Figure S3C). Consequently, HSF1 inactivation reduced heat-shock protein expression such as Hsp90α and Hsp27 (Figure 3F). Moreover, HSP90 clients including c-Raf, AKT and Bcl-xl also destabilized, confirming the inactivation of the HSF1-HSP90 anti-proteotoxic defense response upon p53 activation (Figure 3F).

To further strengthen the evidence for this repressive WTp53-HSF1 axis, we generated stable HSF1-overexpressing HCT116 clones, functionally confirmed by increased levels of pSer326-HSF1 (Figure 3G) and higher expression of HSF1 target genes (Figure S3F). Again, Nutlin strongly dephosphorylated pSer326-HSF1 (Figure 3G) and down-regulated the increased HSF1 target gene response in these clones (Figure S3F). To finally demonstrate that HSF1 is directly controlled by WTp53, we used heat-shock, the strongest known HSF1 activator, to massively increase endogenous HSF1 activity in human CRC cells. Again, Nutlin strongly repressed HSF1 activity, demonstrating how potently WTp53 counter-regulates even the strongest HSF1 activator (Figures 3H-I). In sum, the HSF1-mediated stress response is strongly attenuated by activation of WTp53.

### p53 blocks HSF1 activity via p21-mediated cell cycle inhibition in human CRC cells

To gain further insight into Nutlin-induced HSF1 repression, we analyzed *CDKN1A*/p21, a key p53 target gene and potent cyclin-dependent kinase inhibitor that mediates cell cycle arrest (Figure S4A). Indeed, p21 depletion by siRNAs abolished pSer326-HSF1 dephosphorylation (Figure 4A) and nearly reversed Nutlin-induced HSF1 target gene repression (Figures 4B, S4B), indicating a p53-p21-mediated HSF1 suppression.

**Figure 4.**
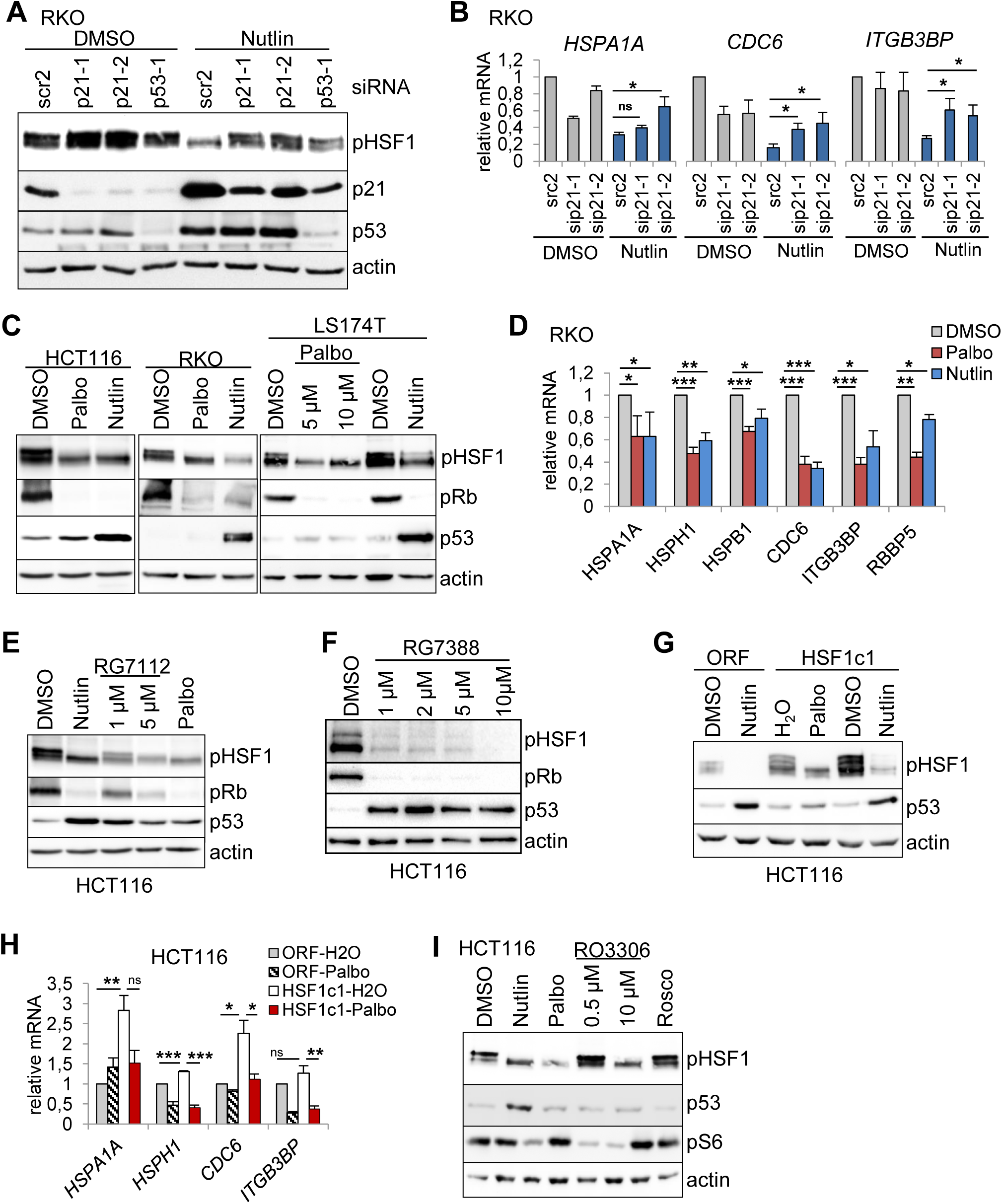
p53 suppresses HSF1 activity via cyclin-dependent kinase inhibitor CDKN1A/p21 in human CRC cells. (A) p21 silencing attenuates p53-induced HSF1 inactivation. RKO cells were transfected with different siRNAs against p21 and p53 or scrambled control siRNA (scr2) for 48 hrs. Cells were then treated with DMSO or 10 µM Nutlin for 24 hrs. Representative Immunoblot analysis for pSer326-HSF1, p21 and p53. Actin, loading control. (B) Rescue of p53-induced HSF1 target gene suppression by depletion of p21 in RKO cells. 48 hrs post transfection with siRNAs against p21 or scrambled control siRNA (scr2), cells were treated with DMSO or 10 µM Nutlin for 24 hrs. qRT-PCR of the indicated mRNAs, normalized to 36B4 mRNA. Relative values given in [ratio (2^-ddCT^)]. Mean ±SEM of 2 independent experiments, each repeated twice in triplicates. Student’s t-test, p*=0.05; ns, not significant. (C) WTp53 harboring CRC cell lines were treated with DMSO, 10 µM Palbociclib (Palbo) or 10 µM Nutlin for 24 hrs. Cell cycle inhibition was confirmed by Rb de(hypo)phosphorylation. Immunoblot analysis for the indicated proteins. pRb, phospho-Rb. Actin, loading control. (D) HSF1 target gene repression after direct cell cycle inhibition. RKO cells were treated with 10 µM Palbociclib (CDK4/6 inhibitor), 10 µM Nutlin (MDM2 inhibitor) or DMSO for 24 hrs. qRT-PCRs analysis for the indicated mRNAs. Relative values calculated as in (B). Mean ±SEM of 2 independent experiments, each repeated twice in triplicates. Student’s t-test, p*=0.05, p**=0.01, p***=0.001; ns, not significant. (E, F) Cell cycle inhibition by p53 inactivates HSF1 activity. HCT116 cells were treated with DMSO, 10 µM Nutlin, 10 µM Palbociclib, RG7112 (E) or RG7388/Idasanutlin (F) as indicated for 24 hrs. Immunoblot analysis. Actin, loading control. (G) Cell cycle inhibition prevents pSer326-HSF1 activation in stably HSF1-overexpressing HCT116 cells. HSF1c1 or empty vector control (ORF) cells were treated with DMSO, H_2_O, 10 µM Nutlin or 10 µM Palbociclib for 24 hrs. Representative immunoblot analysis. Actin, loading control. (H) Cell cycle inhibition in HSF1-overexpressing HSF1c1 cells strongly repress HSF1 target gene expression. HSF1c1 or ORF control cells were exposed to H_2_O or Palbociclib (10 µM) for 24 hrs. qRT-PCR analysis of the indicated HSF1 target genes. Mean ±SEM of 2 independent experiments, each repeated twice in triplicates. Student’s t-test, p*=0.05, p**=0.01, p***=0.001; ns, not significant. (I) Direct CDK4/6 inhibition drives HSF1 inactivation. HCT116 cells were treated with DMSO, 10 µM Nutlin, 10 µM Palbociclib, 0.5 µM and 10 µM RO3306 and 20 µM Roscovitine (Rosco) for 24 hrs. Representative immunoblot. Actin, loading control.

Next we asked whether the p53/p21-mediated HSF1 suppression is linked to and regulated by the cell cycle. *CDKN1A*/p21 binds and inhibits cyclin-dependent kinases (CDKs), thereby preventing phosphorylation of the retinoblastoma protein (RB). Hypo-phosphorylated RB binds to and inhibits E2F transcription factors preventing S-phase entry^59^. Thus, we tested whether cell cycle inhibitors like CDK4/6 inhibitor Palbociclib phenocopy the p53-p21-mediated HSF1 inactivation. Indeed, both Nutlin- and Palbociclib-treated cells exhibited markedly decreased levels of pSer326-HSF1 in WTp53 cells (Figure 4C). Moreover, HSF1 targets were suppressed by Palbociclib, mimicking the Nutlin-induced HSF1 response (Figure 4D). In further support, Nutlin-derivatives RG7112 and RG7388 (Idasanutlin) also reduced pSer326-HSF1 (Figures 4E-F). Likewise, in HSF1-overexpressing HCT116 clones cell cycle inhibition by Palbociclib (like Nutlin) repressed pSer326-HSF1 levels (Figure 4G) and target gene expression (Figure 4H).

To pinpoint the specific CDKs involved in activating HSF1, we used RO3306 (inhibits CDK1 and CDK2 at lower concentrations but CDK4 at higher concentrations) and Roscovitine (inhibits CDK1, CDK2, CDK5 and CDK7, but poorly CDK4/CDK6). Of note, only RO3306 at higher concentrations blocked pSer326-HSF1 like Nutlin and Palbociclib did (Figure 4I), indicating a specific role for CDK4/6 in HSF1 activation. Overall, these data demonstrate that cell cycle inhibition via p53-induced p21-CDK4/6 signaling suppresses HSF1 activity.

### WTp53 activation represses MLK3. MLK3 links cell cycle to the MAPK stress pathway to activate the HSF1 response

The HSF1 stress response is markedly attenuated by CDK4/6 inhibition (Figure 4). Thus, we tested whether E2F target genes like CDK1, CDK2, CDC25C, PLK4 and MLK3 control the HSF1-mediated HSR. Indeed, these E2F targets are all strongly repressed by Nutlin-activated p53, an effect largely rescued by concomitant p53 depletion (Figure 5A). Specifically, MLK3 depletion mimicked the Nutlin response and reduced both pSer326-HSF1 (Figure 5B) and HSF1 target gene expression (Figures 5C-D). Notably, MLK3 directly signals to the MEK/ERK stress pathway^60-62^ and MEK/ERK activates HSF1 by phosphorylation^35, 63, 64^. In contrast, depletion of CDK1 or CDK2 failed to reduce pSer-325 HSF1 (Figures S5A-B). Moreover, PLK4 protein was not diminished after silencing, albeit PLK4 mRNA was strongly reduced, pointing to a stable PLK4 protein but excluding PLK4 as HSF1-activating kinase (Figures S5C, D). In contrast, MLK3 depletion reduced both pSer326-HSF1 and MEK phosphorylation (Figure 5B), revealing a MLK3-MEK-HSF1 signaling axis.

**Figure 5.**
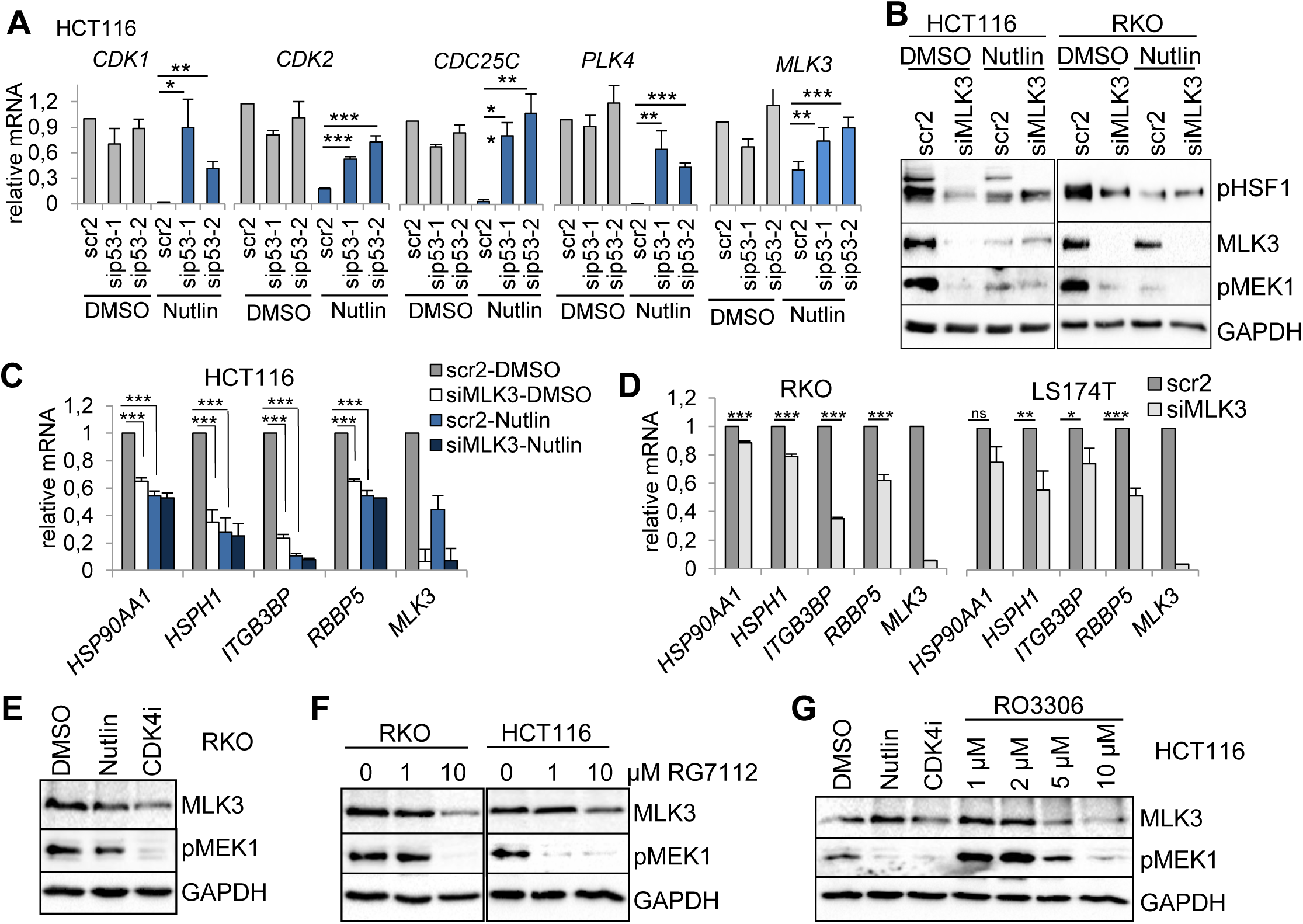
WTp53 activation represses MLK3. MLK3 links cell cycle to the MAPK stress pathway to activate the HSF1 response. (A) Expression of cell cycle progression genes is inhibited by p53 activation. HCT116 cells were transfected with siRNAs for p53 or scrambled control siRNA (scr2) for 48 hrs, followed by DMSO or 10 µM Nutlin treatment for 24 hrs. qRT-PCRs for the indicated mRNAs, each normalized to 36B4 mRNA. MLK3, Mixed lineage kinase 3. PLK4, polo-like kinas 4. Relative values as ratio (2^-ddCT^). Mean ±SEM of 2 independent experiments, each repeated in triplicates. Student’s t-test, p*=0.05, p**=0.01, p***=0.001. (B) MLK3 silencing suppresses pSer326-HSF1, mimicking that seen by p53 activation. Indicated cells were transfected with an siRNAs pool against MLK3 or scrambled control siRNA (scr2). 48 hrs post-transfection, cells were treated with DMSO or 10 µM Nutlin for 24 hrs. Immunoblot analysis. GAPDH, loading control. (C) MLK3 silencing abrogates HSF1 target gene expression, mimicking that seen by p53 activation. HCT116 cells were transfected and treated as in (B). qRT-PCRs for the indicated mRNAs, each normalized to 36B4 mRNA. Relative values were calculated as in (A). Mean ±SEM of 2 independent experiments, each repeated in triplicates. Student’s t-test, p*=0.05, p**=0.01, p***=0.001. (D) HSF1 target gene expression is attenuated after MLK3 depletion. Indicated cells were transfected with an siRNAs pool against MLK3 or scrambled control siRNA (scr2). 72 hrs post-transfection, qRT-PCRs for the indicated mRNAs was performed. Normalized to 36B4 mRNA. Relative values and means as in (A). Mean ±SEM of 2 independent experiments, each repeated in triplicates. Student’s t-test, p*=0.05, p**=0.01, p***=0.001. (E-G) Cell cycle inhibition reduces MLK3 expression and causes MEK1 inactivation. The indicated cells were treated for 24 hrs with DMSO, 10 µM Nutlin or 10 µM Palbociclib (CDK4i) (E, G), RG7112 (F) and RO3306 (G) at the indicated concentrations. Immunoblot analysis. GAPDH, loading control.

Importantly, MLK3 mRNA and protein levels were reduced after p53 activation (Nutlin and RG7112, Figures 5A-B, E-F, S5E) and cell cycle inhibition (CDK4i and RO3306, Figures 5E-G), concomitant with MEK inactivation (pMEK1, Figures 5E-G). Thus, we confirmed the MAPK pathway as major HSF1 activator in human WTp53 harboring CRC cells. Taken together, we identified MLK3 as upstream link between cell cycle and the MAPK stress pathway to activate HSF1. WTp53 activation represses MLK3, which in turn inactivates the MAPK stress pathway and consequently the HSF1 response.

### In human colorectal cancer p53LOH combined with missense mutp53 tends to shorten patient survival and upregulate HSF1 activity

Tumors strongly depend on constitutively upregulated chaperones to manage pervasive proteotoxic stress. However, functional WTp53 prevents adaptive upregulation of the HSF1 chaperone system (Figures 2-5). Thus, to survive and progress, tumors are under strong selection pressure to undergo p53LOH^65^ and lose WTp53-mediated HSF1 repression. This scenario was confirmed in human CRC. p53LOH occurred in ∼80% of patients harboring all p53 variants or missense-only (MS) (Figure 6A, COADREAD TCGA data). Importantly, p53LOH combined with missense mutp53 showed a trend to shorter survival (median 57.2 month *vs*. 83.2 months in WTp53 patients) (Figure 6B, note that TCGA lacks sufficient numbers of heterozygous patients (mutp53/+), precluding statistical analysis). Remarkably, HSF1 target genes are concomitantly upregulated in p53LOH CRCs harboring all p53 mutations (Figure S6A) or missense-only variants (Figure 6C). Moreover, p53LOH breast cancers (BRCA TCGA) also exhibited upregulated HSF1 targets (Figures S6B-C). Furthermore, BRCA cells repressed pSer326-HSF1 when harboring WTp53 but not mutp53 (Figure S6D). In sum, these data support that in human colorectal and breast cancers p53LOH overrides HSF1 repression by WTp53 and enables pleiotropic tumor-promoting HSF1 functions contributing to poorer prognosis.

**Figure 6.**
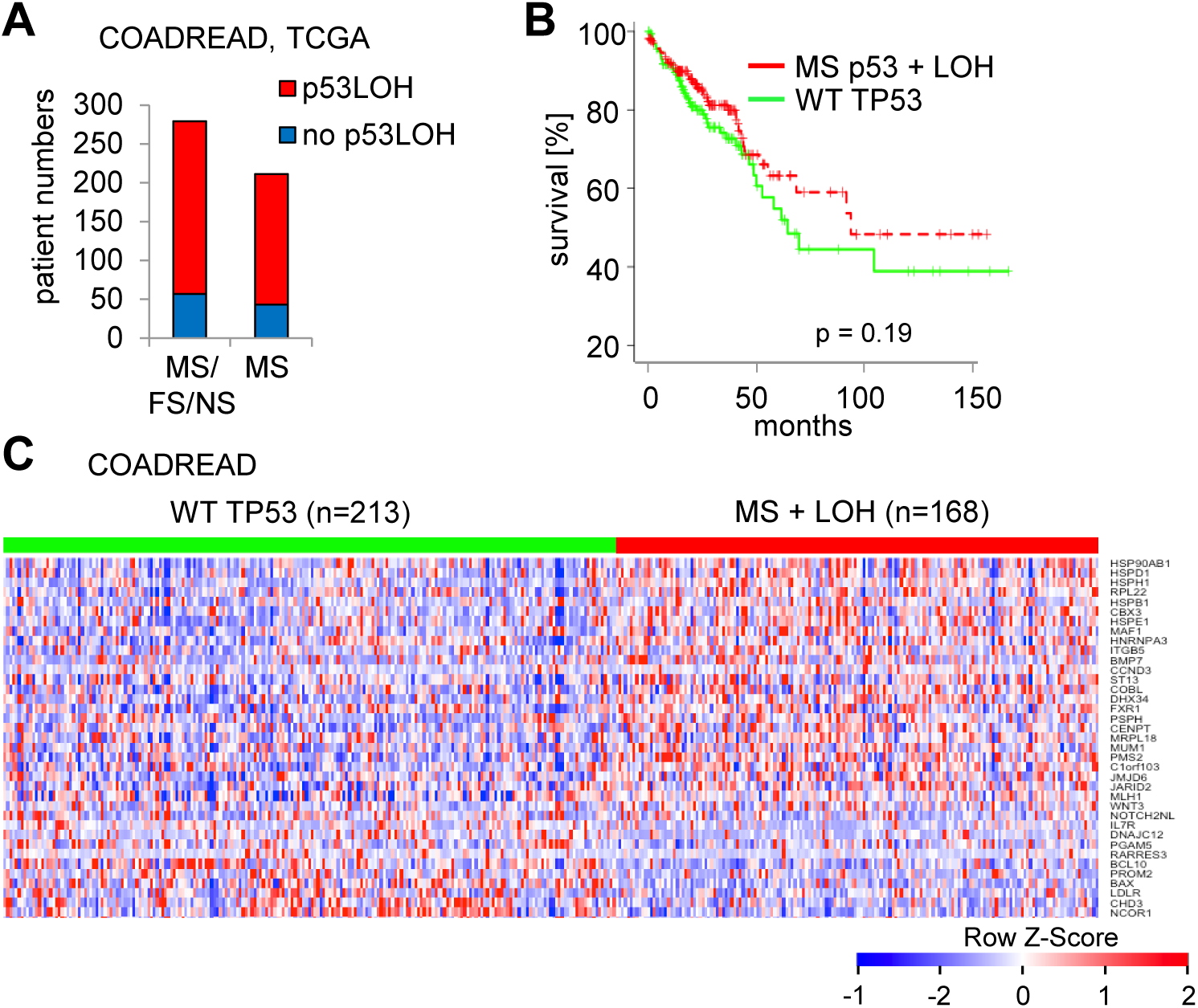
p53LOH combined with p53 missense mutations shortens patient survival and upregulates HSF1 activity in human colorectal cancer. (A) Proportion of human colorectal adenocarcinoma samples with p53LOH versus no-p53LOH. Analysis of the latest version of the COADREAD TCGA dataset, grouped for all types of TP53 mutations (MS, missense; FS, frameshift; NS, nonsense) versus TP53 missense mutations-only (MS). p53LOH samples (= shallow deletion) were determined by TP53 copy number alterations. MS/FS/NS column, p53LOH samples in red of n=222 patients, and no-p53LOH in blue of n=57 patients. MS column, p53LOH samples in red with n=168 patients, and no-p53LOH in blue with n=43. (B) Kaplan-Meier survival curve of all available patients from COADREAD TCGA database. Colorectal cancer patients harboring homozygous WT TP53 were compared to patients harboring missense (MS) p53 mutations plus p53LOH (= shallow deletions). The mean survival of WTp53 patients (n=214) is 83.2 month versus 57.2 months for patients with MS plus p53LOH (mutp53/-) (n=166). Kaplan-Meier statistic on patient cohorts from TCGA, log-rank test, p=0.19. Note that TCGA data contains insufficient numbers of heterozygous patients (mutp53/+), precluding statistical analysis. (C) Heatmap of HSF1 target genes analyzed from colorectal adenocarcinoma patients (COADREAD cohort TCGA database). Patients harboring homozygous TP53^+/+^ (WT TP53) were compared to patients harboring TP53 missense (MS) mutations plus p53LOH (= shallow deletions) (TP53^MS + LOH^). Patient numbers are indicated. Genes were ordered from top to bottom by their relative upregulation (red) and downregulation (blue) and their p-value significance in t-tests. The HSF1 target gene panel from Mendillo et al was used^39^. Note, HSF1 negatively regulates a subset of target genes.

### In murine CRC organoids p53LOH enables HSF1 activity and triggers mutp53 stabilization

To further strengthen the WTp53-induced HSF1 repression in heterozygous epithelial cells, we generated tumor-derived organoids from our murine CRCs (Figure 7A). Importantly, p53^Q/fl^ organoid cultures maintain their heterozygous p53 status over at least 8 passages (Figures S7A-B). Thus, we treated p53^Q/fl^; vilCreER^T2^ tumor-derived organoids first with 4OHT to induce p53LOH, followed by Nutlin (Figures 7A-B). Heterozygous organoids (EtOH controls) showed strong induction of p53 target genes after Nutlin, whereas the p53 response was significantly damped in p53LOH organoids (4OHT group) (Figure 7B). Cre recombinase-mediated allele deletion is never 100% efficient, creating competition between non-recombined and recombined tumor cells. Indeed, p53 mRNA levels post 4OHT were above those corresponding to a single (mutant) TP53 copy (Figure S7C), indicating that recombination was below 100%, and explaining the mild but detectable Nutlin response in 4OHT-treated organoids (Figure 7B). Importantly, HSF1 target genes were de-repressed in p53LOH organoids (4OHT group) after Nutlin versus the no-LOH (EtOH) group (Figure 7C), again confirming that p53LOH enables HSF1 activity.

**Figure 7.**
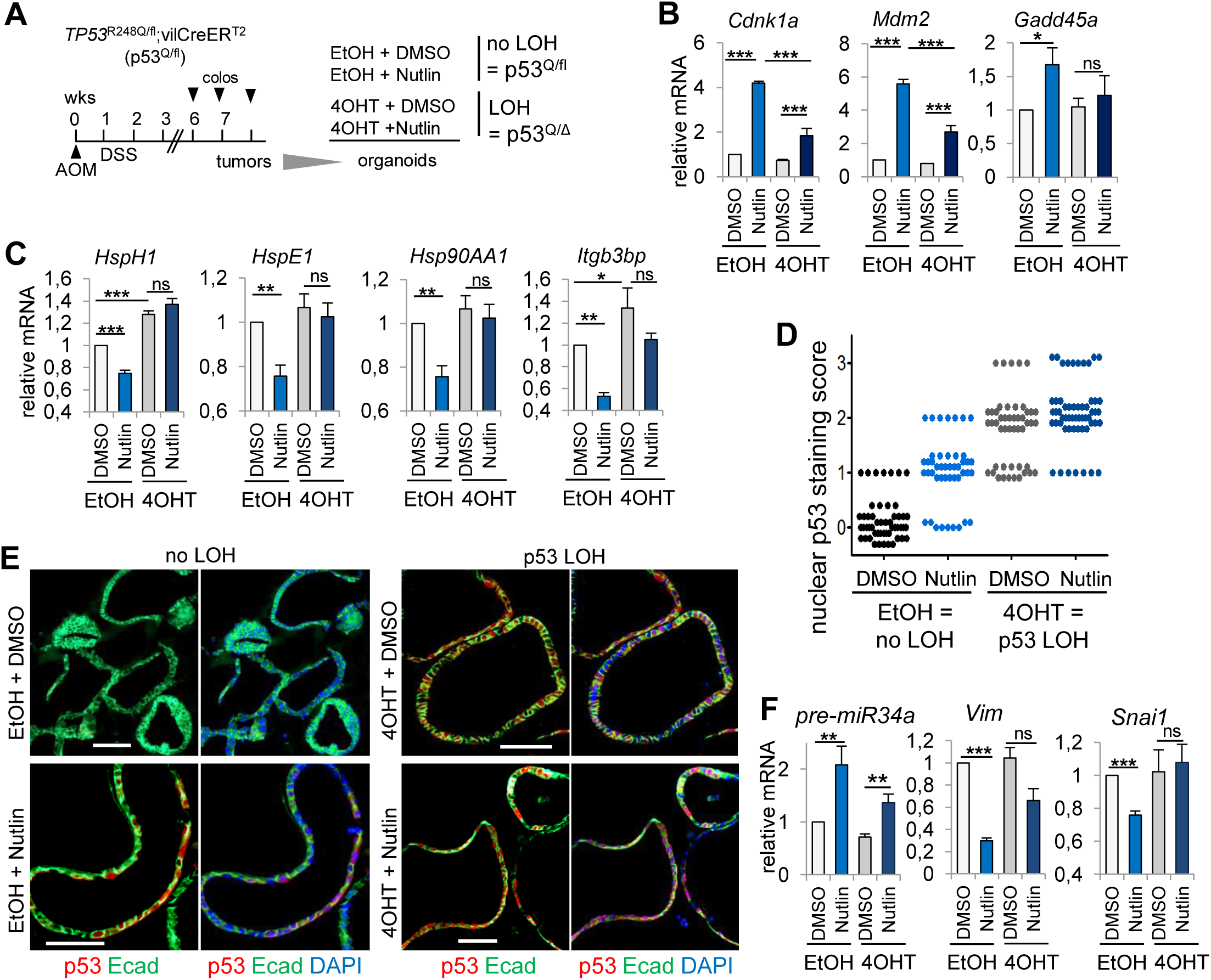
In murine CRC organoids p53LOH enables HSF1 activity and triggers mutp53 stabilization. (A) Scheme for treatment of colonic tumor-derived organoids. Heterozygous p53^Q/fl^; vilCreER^T2^ mice were treated with AOM/DSS and tumor burden was visualized via colonoscopy. Tumors arisen between 6-8 wks post AOM were resected and processed for colonic organoid cultures. p53LOH was induced by adding 4OHT (4OH-Tamoxifen) for 24 hrs to activate the CreER^T2^ recombinase and create p53^Q/Δ^ organoids. EtOH, control treatment (no-LOH, p53^Q/fl^). Two days after p53LOH induction, organoids were treated with 10 µM Nutlin or DMSO for 24 hrs and harvested for analysis. (B, C) mRNA levels of p53 target genes (B) and HSF1 target genes (C) isolated from colonic p53^Q/fl^ organoids treated as indicated. qRT-PCR normalized to HPRT mRNA. Mean ± SEM of 3 different organoid cultures (generated from 3 different mice) each measured in triplicates. Student’s t-test, p*=0.05, p**=0.01, p***=0.001; ns, not significant. (D) Quantification of (E) using a score for colonic organoids exhibiting nuclear p53 staining. Each dot indicates one organoid of the indicated treatment groups. Nuclear p53 staining score: 0 = no positive nucleus per organoid; 1 = 1 – 20% positive nuclei per organoid; 2 = 20 – 50% positive nuclei per organoid and 3 > 50 % positive nuclei per organoid. (E) Representative immunofluorescence staining of the indicated p53^Q/fl^ organoid groups for p53 (red), E-cadherin (Ecad, green) and DAPI (blue). Scale bars, 100 μm. (F) mRNA levels of pre-miR34a, Vim and Snai1 of colonic p53^Q/fl^ organoids treated as indicated. qRT-PCR normalized to HPRT mRNA. Mean ± SEM of 3 different organoid cultures (generated from 3 different mice) each measured in triplicates. Student’s t-test, p**=0.01, p***=0.001; ns, not significant.

In agreement with an upregulated chaperone system after p53LOH, nuclear mutp53^R248Q^ became strongly stabilized after 4OHT (Figures 7D-E). Moreover, while WTp53 strongly prevents invasion by upregulating e.g. miR34a which controls epithelial-to-mesenchymal transition (EMT)^66^, miR34a induction is markedly diminished after p53LOH, allowing upregulation of EMT markers Vimentin and SnaiI (Figure 7F, compare EtOH+Nutlin *vs* 4OHT+Nutlin). Combined upregulation of EMT genes and pro-invasive HSF1 target genes like HspH1^67^ and Itgb3bp^39^ (Figure 7C) enable invasiveness in a stressed tumor milieu following p53LOH.

## DISCUSSION

Here we use an autochthonous immune-competent mouse model that recapitulates human colorectal cancer^68^ and identify the mechanism that drives the critical p53LOH event in p53 missense mutant heterozygous tumors. We show that the GOF p53^R248Q^ allele fails to exert a dominant-negative effect over the remaining WTp53 allele. Instead, the activated WT allele retains its tumor-suppressive function and strongly suppresses the potent HSF1-mediated stress response necessary for tumor maintenance and progression. We identify a repressive WTp53-p21-MLK3-MAPK-HSF1 signaling cascade as the underlying mechanism that creates the driving force for losing the WTp53 allele. The retained WTp53 allele, via its ability to repress the HSF1-regulated chaperone system, prevents stabilization of mutp53 protein in heterozygous tumors, thereby blocking the full oncogenic potency of GOF alleles. Hence, WTp53-mediated HSF1 suppression exerts a strong selection pressure for p53LOH. Conversely, p53LOH, once it occurs, is a dramatic all-or-none gate-opener, unleashing the broad GOF functions of the mutant allele by mediating mutp53 stabilization via upregulating the HSF1-HSP90 chaperone system. This in turn enables tumor invasion^8, 9, 16, 69^. Our findings, corroborated by CRC murine organoids, human cell lines and human tumors in which mutp53 enables the critical transition from late adenoma to invasive carcinoma^1, 2^, reveal the pivotal significance of the repressive WTp53-HSF1 axis. Thus, a single genetic event, p53LOH, kills 3 birds with one stone: *i*) losing WTp53 suppressor activity including HSF1 repression, *ii)* upregulating tumor-promoting HSF1, and *iii*) enabling mutp53 protein stabilization, thereby unleashing the GOF potential.

Studies of normal tissues of mutp53 knockin mice established that MDM2 degrades mutant and WTp53 equally well, keeping both mutant and WT levels below immunohistochemical detection^16, 18, 42^. Conversely, in response to stress both WT and mutp53 proteins stabilize^56, 70, 71^. Importantly, we find in our CRC model that the remaining WT allele in heterozygous tumors is fully activatable, excluding a dominant-negative effect (DNE) by the counterpart mutp53 allele. Although DNE is a likely key driver behind the p53 mutational spectrum in myeloid malignancies^72^, many missense mutants are highly inefficient in their ability to exert DNE in epithelial carcinomas and sarcomas, including the hotspot p53^R248Q^ allele^9, 31, 56, 73^. We and others speculate that for DNE to occur in solid tumors, the MUT/WT protein ratio has to greatly shift in favor of MUT^56, 74^. Yet, while DNE requires acute stress to increase the low-abundance mutp53 protein and stoichiometrically overwhelm co-expressed WTp53, stress equally stabilizes WTp53 levels, thus not shifting the ratio. Only a highly active HSF1-chaperone system with its mutp53-selective accumulation can induce the required MUT predominance over WT. However, the repressive WTp53-HSF1 axis prevents such unilateral mutp53 stabilization, explaining the strong selection pressure for p53LOH.

AOM/DSS CRCs require WTp53 activation by Nutlin (mimicking high proliferative stress or chemotherapy) to fully regulate HSF1 (Figures 2D, S2G), patient p53LOH tumors intrinsically exhibit upregulated HSF1 targets compared to WTp53 tumors (Figures 6C, S6A-C). We speculate that heterozygous human tumors are sufficiently *constitutively* stressed to activate WTp53 and repress HSF1, and that upon p53LOH human tumors massively upregulate HSF1 activity. In contrast, we posit that baseline AOM/DSS-induced tumors have insufficient stress levels to drive p53LOH spontaneously, explaining the missing spontaneous p53LOH in the AOM/DSS model (Figures 1F, S1B), compared to KRAS-driven mouse models of pancreas and lung cancer^42, 45^. Notably, in human CRCs strong constitutive oncogenic stress from K-RAS/ EGFR/ TGFβR/ PDGFR mutations are preeminent^65, 69, 75, 76^, while AOM/DSS tumors undergo predominantly CTNNB1 but no K-Ras and other proliferative driver mutations^68, 77^ which promote proliferative stress^65^. Thus, baseline murine CRCs might not be stressed enough to spontaneously activate WTp53 and suppress HSF1 (Figure S2G). This might also explain the preferred order in which cancer-causing mutations occur during tumorigenesis^76, 78^.

Heat shock-induced accumulation and activation of WTp53 also depends on HSF1^40, 79, 80^ and HSF1/chaperone functions^71, 81-84^. Here we identify a hitherto unknown repressive feedback WTp53-HSF1 counter-regulation. Our data provide an explanation for the longstanding puzzle why tumor heterozygosity tends to be unstable and why p53LOH strictly correlates with mutp53 stabilization and higher tumor aggressiveness.

## Acknowledgement

We thank Nina Pfisterer and Lukas Gebauer (Molecular Medicine MSc-Program Göttingen) for technical assistance. R.S.-H. is supported by the DFG (SCHUH3160/3-1), the Heidenreich-von-Siebold Program (University Medical Center Göttingen) and the KH-Bauer Program (G-CCC Göttingen). U.M.M. is supported by NIH NCI (2R01CA176647), Deutsche Forschungsgemeinschaft (MO1998/2-1) and the Stony Brook Foundation TRO program.

## Figure Legends

**Supplemental Figure 1.**
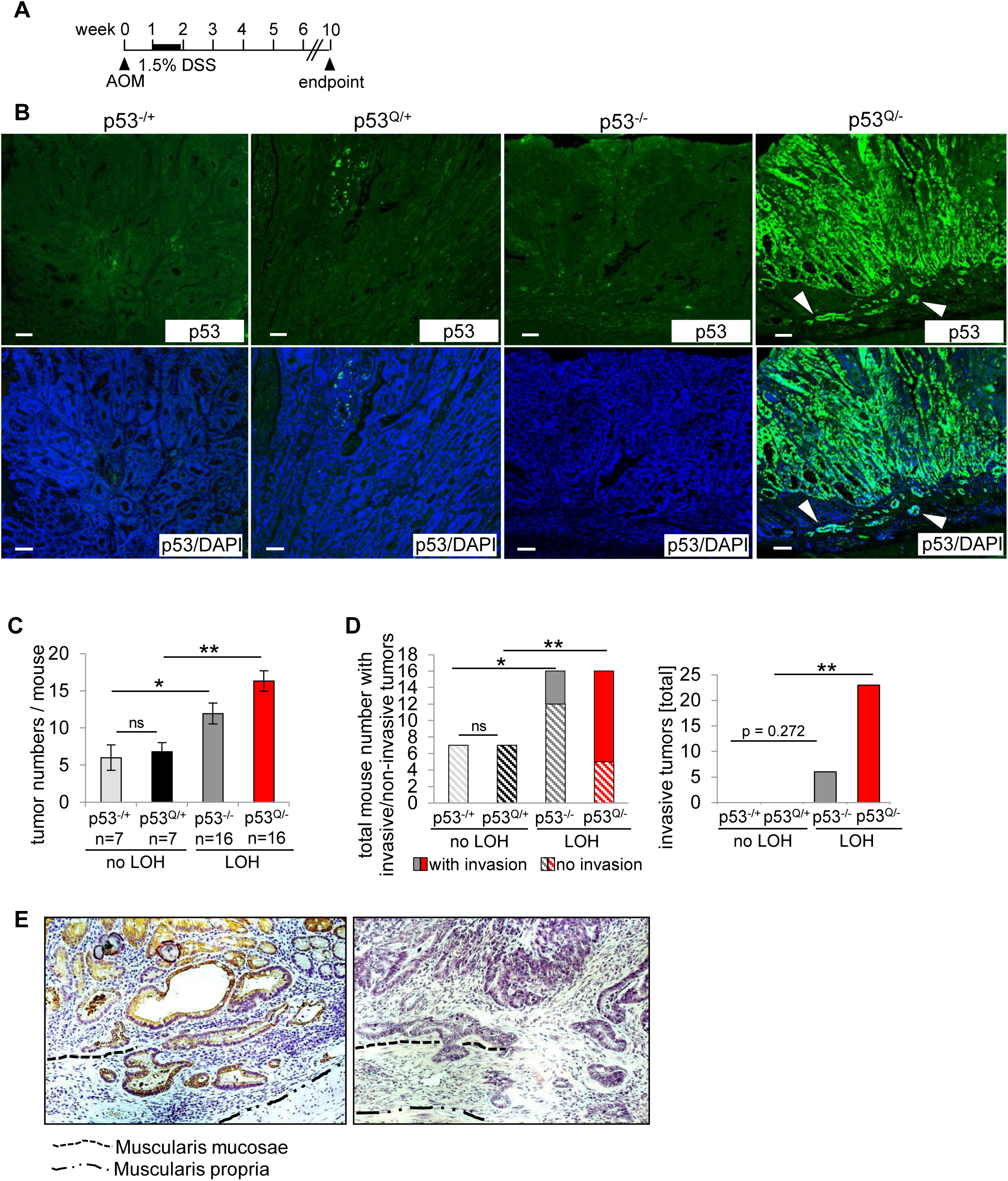
p53 loss-of-heterozygosity is a prerequisite for mutp53 protein stabilization and enables invasion in colorectal cancer. (A) The humanized GOF TP53^R248Q^ allele (p53^Q^) was paired with the p53null allele^85^ in the AOM/DSS colorectal cancer model previously described in Schulz-Heddergott et al.^9^ to generate heterozygous p53^Q/+^ mice (mimicking no LOH) and GOF p53^Q/-^ mice (mimicking p53LOH), with corresponding controls (p53^-/+^ and p53^-/-^ mice). All mice were treated with 1.5% DSS. Time line for p53-proficient (containing one WTp53 allele) and p53-deficient (both p53 alleles are altered) mice used in this study. Endpoint analysis at 10 wks for all mice to avoid losing p53-deficient mice due to lymphoma and intestinal obstruction. (B) Representative immunofluorescence staining for p53 (green) and DAPI (blue) of CRC tumors from the indicated genotypes at endpoint 10 wks. Occasional p53^Q/+^ tumors show a minor focus of stabilized mutp53, presumably an area that underwent p53LOH. Scale bars, 100 μm. White arrowheads show invasive malignant glands. (C) Total tumor numbers per mouse of the indicated genotypes at endpoints described in (A). p53^-/+^ and p53^Q/+^ mice harbor heterozygous CRC tumors. Tumors from p53^-/-^ and p53^Q/-^ mice are homozygous for their TP53 alteration mimicking p53LOH. Bars indicate mean ± SEM, Student’s t-test. p*=0.05; p***=0.001; ns, not significant. (D) Total number of mice with non-invasive and invasive tumors (left) and total number of invasive tumors (right) of the indicated genotypes from (A-C) at endpoints described in (A). (left) p53^-/+^ and p53^Q/+^, n = 7 mice each. p53^-/-^ and p53^Q/-^, n = 16 mice each. (right) p53^-/+^, n = 42 tumors from 7 mice; p53^Q/+^, n = 45 tumors from 7 mice; p53^-/-^, n = 71 tumors from 16 mice and p53^Q/-^, n = 115 tumors from 16 mice. Bars, mean ± SEM. Fisher’s exact test. p***≤ 0.05, p**≤ 0.01. (E) Representative histopathology of p53^Q/-^ tumors. H&E staining, Scale bars, 100 μm. Dashed line, muscularis mucosae; dashed/dot line, border to muscularis propra.

**Supplemental Figure 2.**
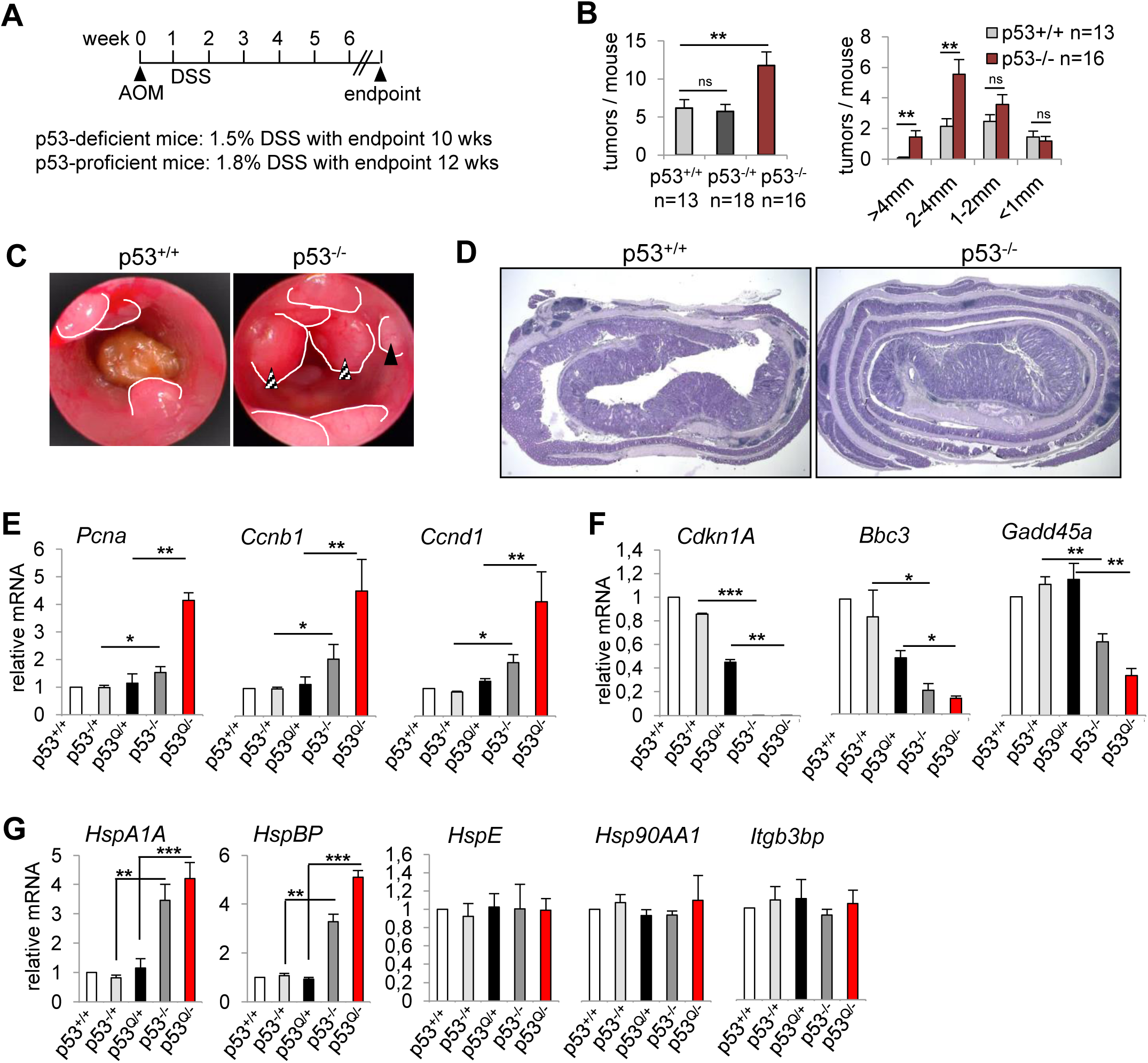
p53 deletion alone is not sufficient to activate HSF1 *in vivo*. (A) Scheme and time line of the AOM/DSS colorectal cancer model using p53null mice^85^. Mice were treated as indicated. Endpoint analysis at 12 wks for p53-proficient mice to avoid losing them to extraneous reasons such as intestinal obstruction and anal prolapse. Endpoint analysis at 10 wks for p53-deficient mice to avoid losing them to lymphoma. (B) Total number of colonic tumors per mouse (left) and tumor size distribution (right) of the indicated genotypes from (A). n, total mouse numbers. Bars, mean ± SEM, Student’s t-test. p**=0.01; ns, not significant. (C) Representative colonoscopy of p53^+/+^ and p53^-/-^ mice at endpoint 10 wks post AOM/DSS. White lines outline tumors. Black arrow indicates an S2 tumor and striped arrows indicate S3 tumors. Tumor scoring was performed according to Becker & Neurath^50^. (D) Colon sections from ‘Swiss roles’ of AOM/DSS-treated p53^+/+^ and p53^-/-^ mice. H&E. (E-G) mRNA levels of cell cycle genes (E), wildtype p53 target genes (F) and HSF1 target genes (G) isolated from the indicated genotypes of colonic tumors (pooled samples, n ≥ 5 tumors per genotype). qRT-PCR normalized to 36B4 or HPRT mRNA. Mean ± SEM of 3 technical replicates, in triplicates. Student’s t-test.

**Supplemental Figure 3.**
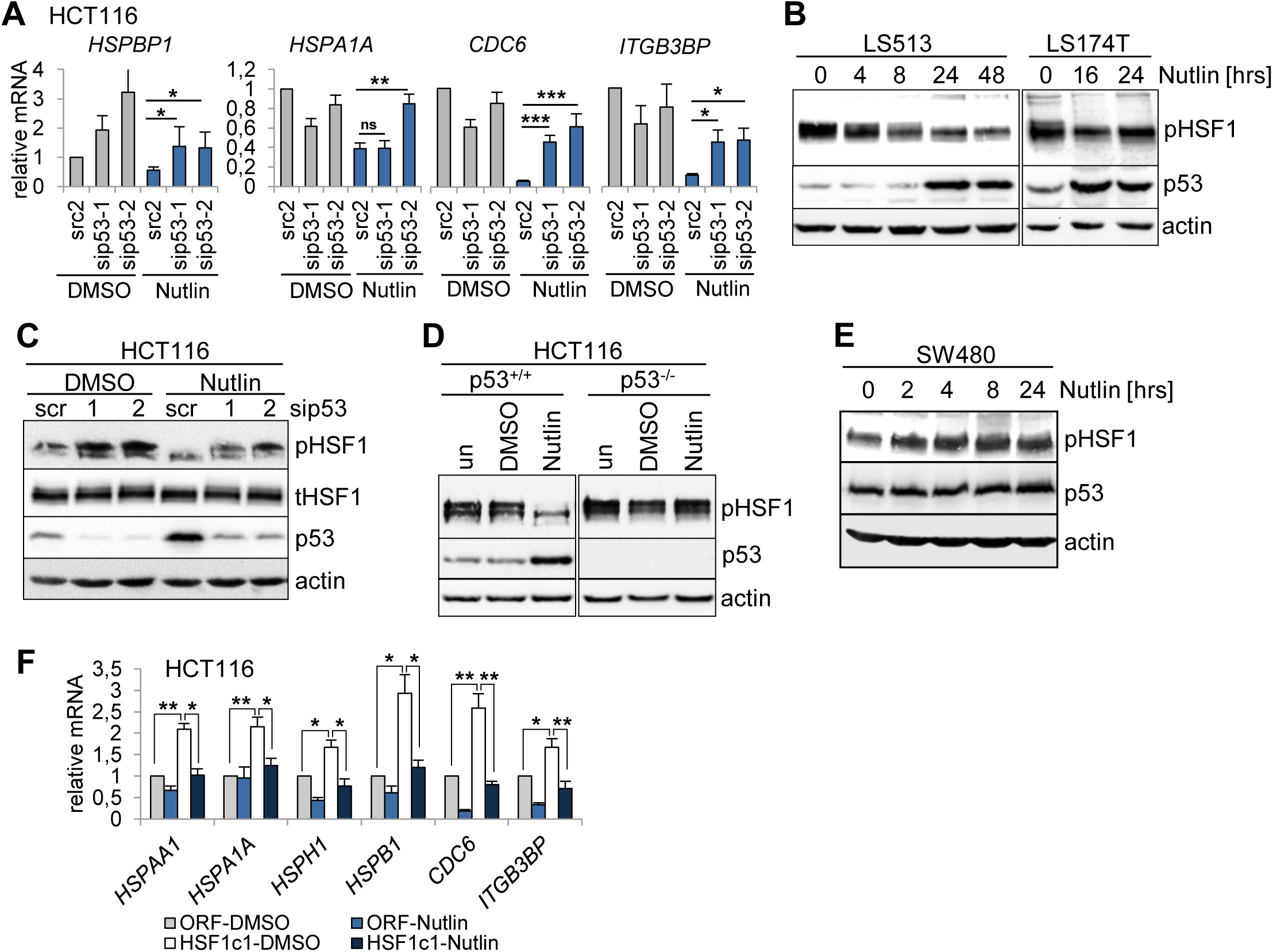
HSF1 activity is repressed by WTp53 in human colorectal cancer cells. (A) p53-induced HSF1 target gene repression is rescued by WTp53 silencing. HCT116 cells were transfected with siRNAs for p53 or scrambled control siRNA (scr2) for 48 hrs. Cells were treated with DMSO or 10 µM Nutlin for 24 hrs. qRT-PCRs for the indicated mRNAs, each normalized to 36B4 mRNA. Relative values are given in [ratio (2^-ddCT^)]. Mean ±SEM of 2 independent experiments, each repeated in triplicates. Student’s t-test, p*=0.05, p**=0.01, p***=0.001; ns, not significant. (B) WTp53 harboring LS513 and LS174T cells were treated with DMSO or 10 µM Nutlin for the indicated times. Representative immunoblot analysis for pSer326-HSF1, the key marker of HSF1 activity. Actin, loading control. (C) p53 silencing abrogates HSF1 inactivation upon Nutlin. HCT116 cells were transfected with two different siRNAs against p53 or scrambled control siRNA (scr). 48 hrs post-transfection, cells were treated with DMSO or 10 µM Nutlin for 24 hrs. Cell lysates were immunoblotted for pSer326-HSF1, total HSF1 (tHSF1) and p53. Actin, loading control. (D) p53 deletion prevents Nutlin-induced HSF1 inactivation. Isogenic HCT116 cells (p53^+/+^ vs p53-/-, harboring a p53 Exon2 deletion) were left untreated (un) or treated with DMSO or Nutlin for 24 hrs. Representative immunoblots for pSer326-HSF1 and p53. Actin, loading control. (E) mutp53-containing CRC cells failed to reduce pSer326-HSF1 after Nutlin. SW480 cells treated +/- Nutlin (20 µM) for the indicated hours. Representative immunoblot. Actin, loading control. (F) Stably HSF1-overexpressing HCT116 subclone HSF1c1 and its empty vector control line (ORF) were treated with DMSO or 10 µM Nutlin for 24 hrs. qRT-PCR analysis of the indicated HSF1 target genes. Mean ±SEM of 2 independent experiments, each repeated twice in triplicates. Student’s t-test, p*=0.05, p**=0.01; ns.

**Supplemental Figure 4.**
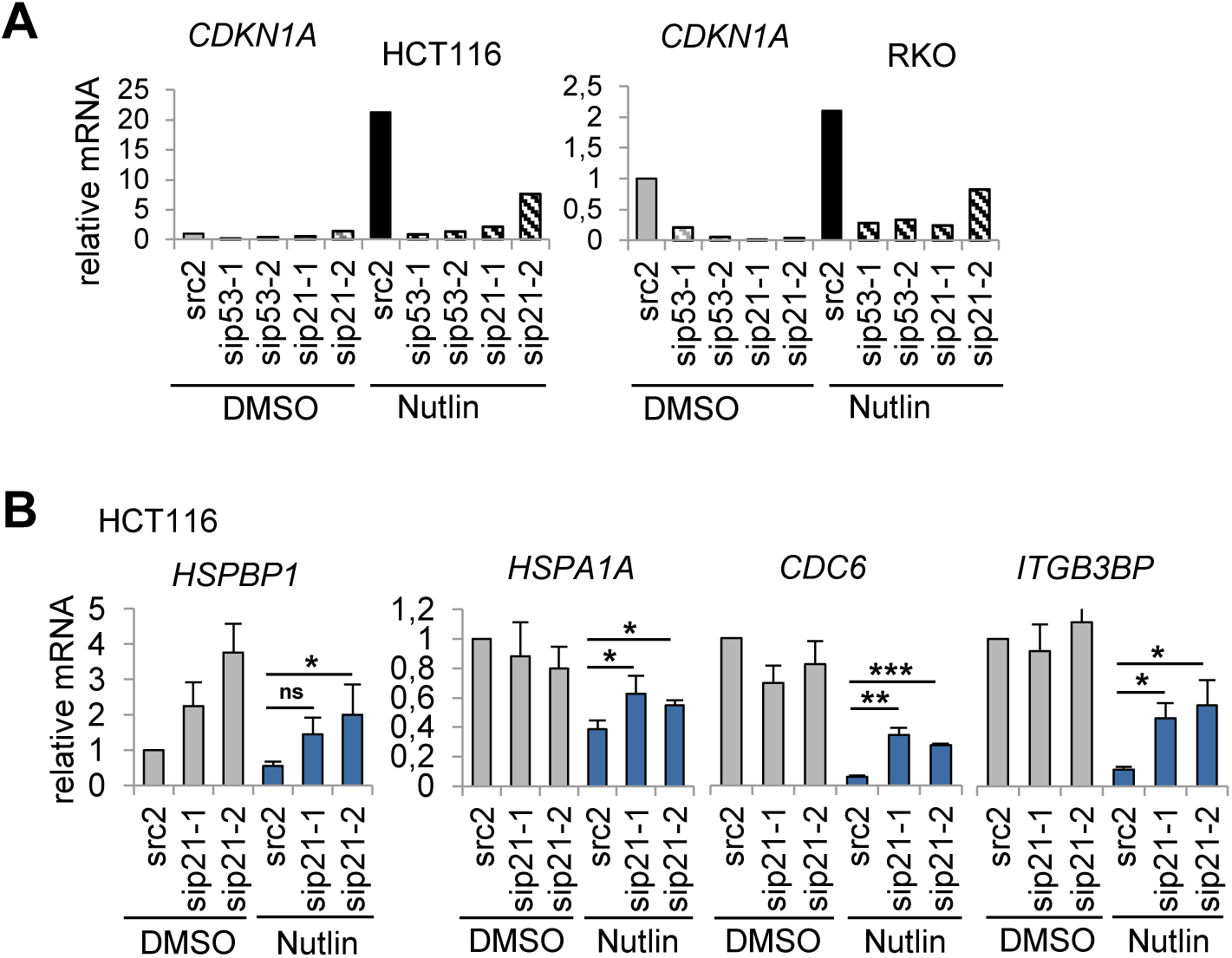
p53 suppresses HSF1 activity via cyclin-dependent kinase inhibitor CDKN1A/p21 in human CRC cells. (A) Analysis of *CDKN1A*/p21 mRNA expression. One representative qRT-PCR each shown for HCT116 and RKO cells. Cells were transfected with siRNAs against *CDKN1A*/p21 and p53 or scrambled control siRNA (scr2) for 48 hrs, followed by DMSO or 10 µM Nutlin treatment for 24 hrs. Relative values of CDKN1A mRNA normalized to 36B4 mRNA and given in [ratio (2^-ddCT^)]. (B) Analysis of HSF1 target gene expression in HCT116 cells upon depletion of p21. 48 hrs post transfection with two siRNAs against p21 or scrambled control siRNA (scr2), HCT116 cells were treated with DMSO or 10 µM Nutlin for 24 hrs. qRT-PCR analysis. Mean ±SEM of 2 independent experiments, each repeated twice in triplicates. Relative values were calculated as in (A). Student’s t-test, p*=0.05, p**=0.01, p***=0.001; ns, not significant.

**Supplemental Figure 5.**
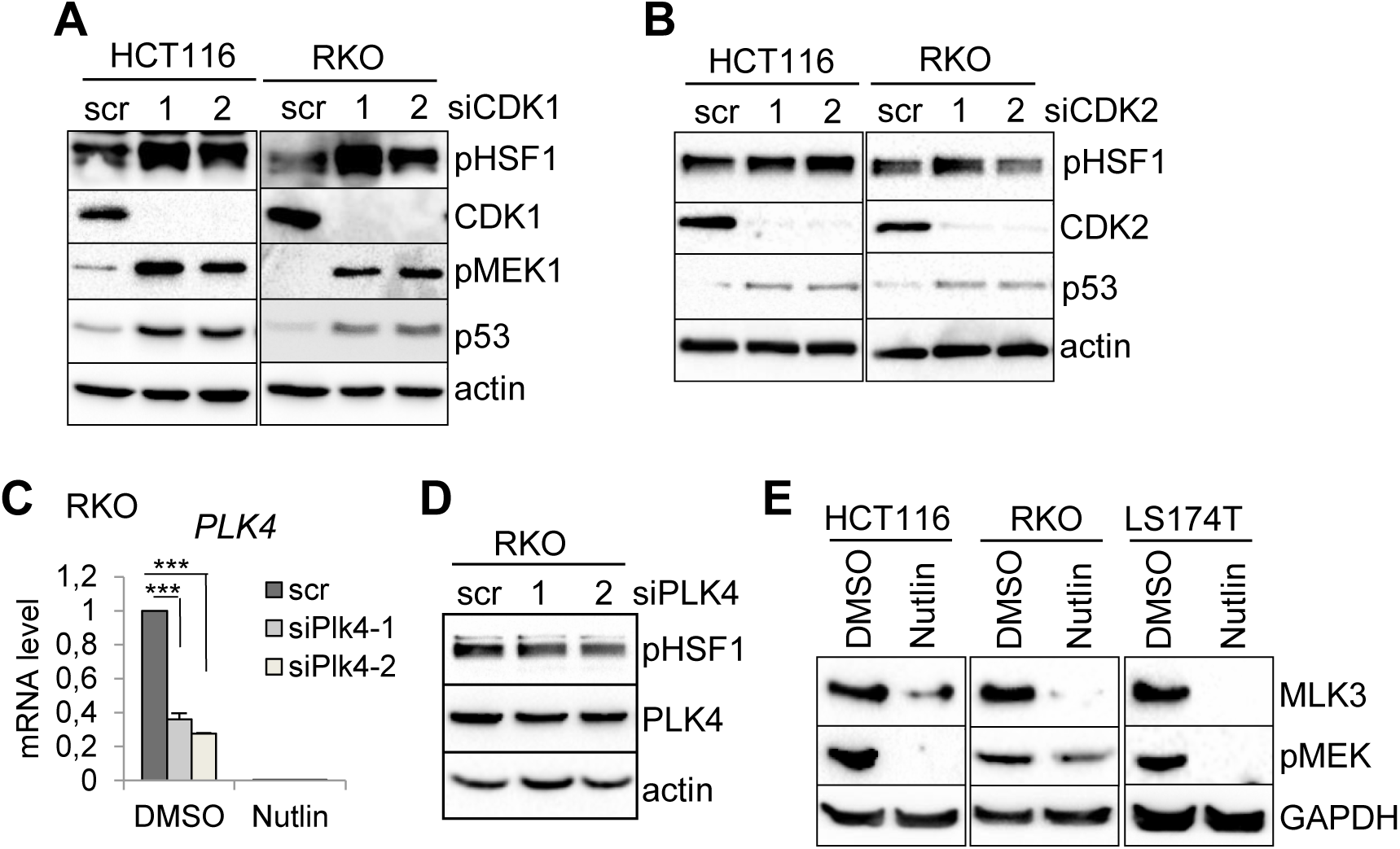
Cell cycle aberrations activate the MEK pathway and regulate HSF1 activity. (A, B) Depletion of CDK1 (A) and CDK2 (B) fail to abrogate HSF1 activity. The indicated cells were transfected with two different siRNAs each. 72 hrs post-transfection, cell lysates were analyzed by immunoblots. Actin, loading control. C) PLK4 silencing in RKO cells from (D). Cells were transfected with 2 different siRNAs against PLK4 or scrambled control siRNA (scr) for 72 hrs. qRT-PCRs for PLK4 mRNAs normalized to 36B4 mRNA. Relative values, ratio (2^-ddCT^). Mean ±SEM of 2 independent experiments, each repeated in triplicates. Student’s t-test, p*=0.05, p**=0.01, p***=0.001. (D) Despite PLK4 mRNA silencing (C), PLK4 protein and pSer326-HSF1 levels are stable, excluding PLK4 as HSF1-activating kinase. Immunoblot analysis of RKO cells from (C). (E) Activated WTp53 strongly reduces MLK3 protein levels. The indicated CRC cells were treated with DMSO or 10 µM Nutlin for 36 hrs. Immunoblot analysis. Actin, loading control.

**Supplemental Figure 6.**
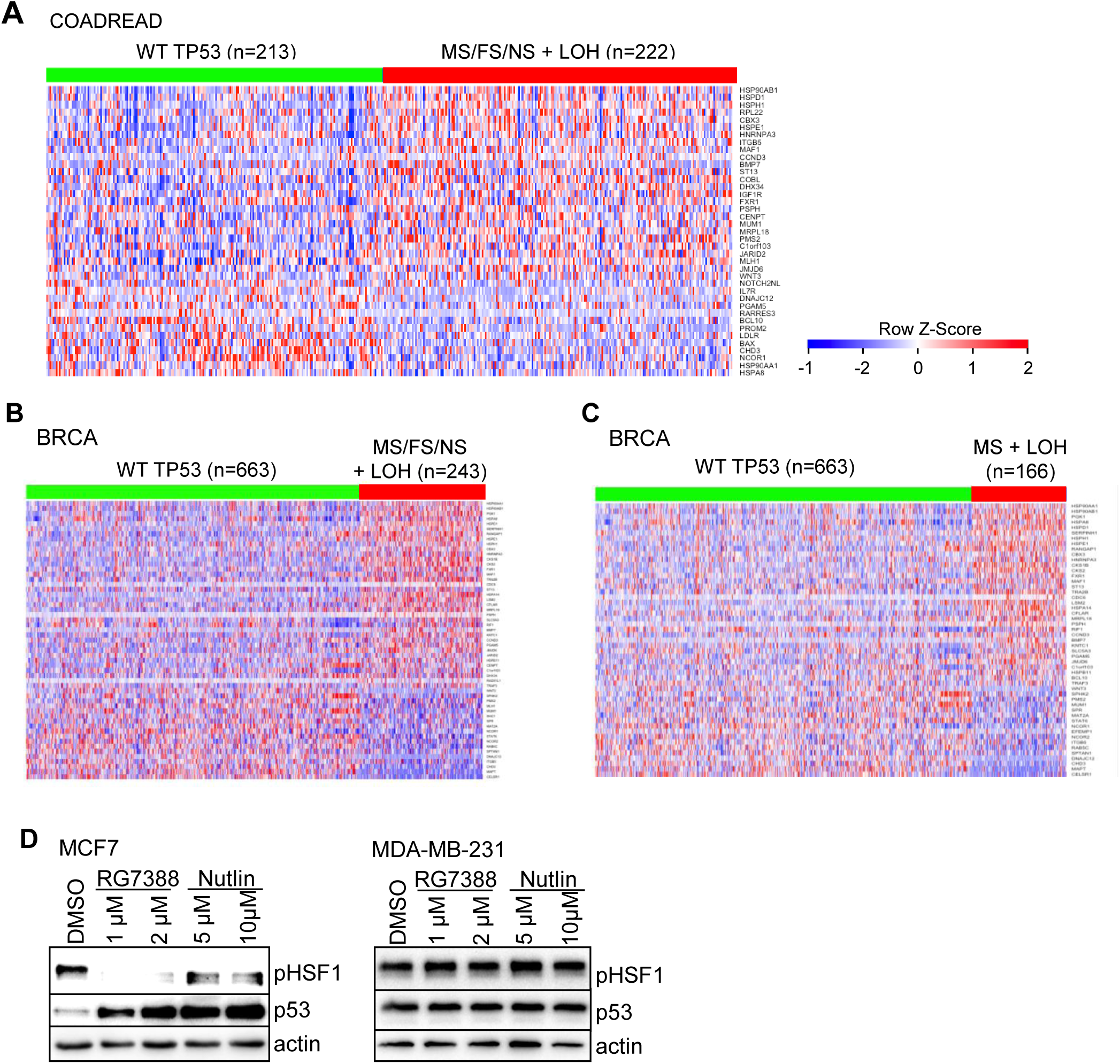
p53LOH combined with p53 missense mutations upregulates HSF1 activity in colorectal and breast cancer patients. (A) Heatmap of HSF1 target genes analyzed from colorectal adenocarcinoma patients (COADREAD cohort TCGA database). Patients harboring homozygous TP53^+/+^ (WT TP53) were compared to patients harboring TP53 alterations (MS, missense; FS, frameshift; NS, nonsense) plus p53LOH (= shallow deletions) (TP53^MS/FS/NS + LOH^). Patient numbers are indicated. (A-C) Genes were ordered from top to bottom by their relative upregulation (red) and downregulation (blue) and their p-value significance in t-tests. The HSF1 target gene panel from Mendillo et al was used^39^. Note, HSF1 negatively regulates a subset of target genes. (B, C) Heatmap of HSF1 target genes analyzed from breast cancer patients (BRCA cohort TCGA database). Patients harboring homozygous TP53^+/+^ (WT TP53) were compared to patients harboring all TP53 alterations (MS, missense; FS, frameshift; NS, nonsense) plus p53LOH (= shallow deletions) (TP53^MS/FS/NS + LOH^) in (B), and also compared to patients harboring TP53 missense (MS) mutations-only plus p53LOH (= shallow deletions) (TP53^MS + LOH^) in (C). Patient numbers are indicated. (D) The MCF7 breast cancer cell line harboring homozygous WTp53 and the MDA-MB-231 breast cancer cell line harboring homozygous mutp53 R280K missense mutation were treated with DMSO, Nutlin or Idasanutlin (RG7388) for 24 hrs as indicated. Representative immunoblot analysis. Actin, loading control.

**Supplemental Figure 7.**
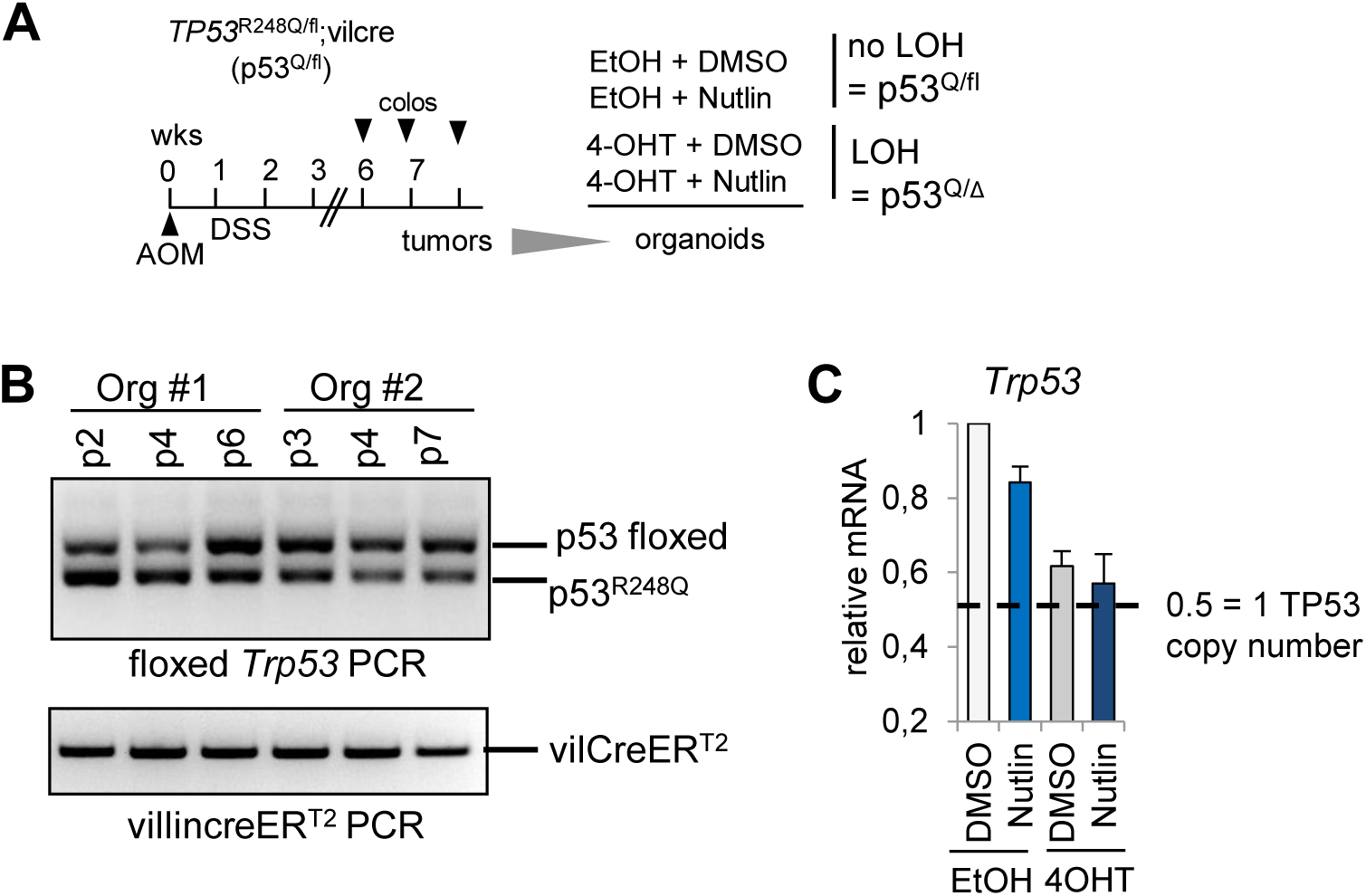
Analysis of heterozygous CRC organoids. (A) Scheme for treatment of colonic tumor-derived organoids. Heterozygous p53^Q/fl^; vilCreER^T2^ mice were treated with AOM/DSS and tumor burden was visualized via colonoscopy. Tumors arisen between 6-8 wks post AOM were resected and processed for colonic organoid cultures. p53LOH was induced by adding 4OHT (4OH-Tamoxifen) for 24 hrs to activate the CreER^T2^ recombinase and create p53^Q/Δ^ organoids. EtOH, control treatment (no-LOH, p53^Q/fl^). Two days after p53LOH induction, organoids were treated with 10 µM Nutlin or DMSO for 24 hrs and harvested for analysis. (B) The heterozygous p53 genotype in CRC organoids is stable. Two randomly chosen organoid cultures (generated from 2 different heterozygous TP53^R248Q/fl^; vilCreER^T2^ mice) were followed during p2-p7 passaging in vitro. The p53 floxed allele and the p53^Q^ allele are indicated. (C) Incomplete recombination. *Trp53* mRNA levels isolated from colonic organoids after p53LOH induction by 4OHT treatment. qRT-PCR normalized to HPRT mRNA. Mean ± SEM of 3 different organoid cultures generated from 3 different mice each measured in triplicates. The dotted line indicates the value corresponding to 1 copy of the TP53 gene.

## Online METHODS

### Mouse experiments and genotyping

Experiments using animal materials were approved by institutional (Göttingen University Medical Center Ethikkommission) and state (Niedersächsisches Landesamt für Verbraucherschutz und Lebensmittelsicherheit, LAVES, Lower Saxony, Germany) committees, ensuring that all experiments conform to the relevant regulatory standards.

The humanized constitutive *TP53*^*R248Q*^ (called p53^Q^) knock-in allele has been described in detail^1-3^. Briefly, the human *TP53* sequence containing the R248Q mutation in exon 7 replaces part of the mouse *Trp53* (exons 4-9). To generate heterozygous mice with one conditional murine Trp53 wildtype allele (p53^fl^), we crossed mice harboring the p53^Q^ allele with mice harboring the floxed WTp53 allele^4^ flanked by loxP sites in introns 2 and 10 to generate p53^Q/fl^. To remove the floxed WTp53 allele from colonic epithelial tissue, we crossed p53^Q/fl^ mice with *villinCreER*^*T2*^ (called ‘ERT2’) transgenic mice.

Moreover, the classic Trp53 knock-out mice (p53^-/-^ mouse)^5^ were crossed to the p53^Q^ allele to generate non-tissue specific TP53 alterations (e.g. Supplemental Figure 1, Supplemental Figure 2) as described in Schulz-Heddergott et al. 2018 ^3^.

For all genotypings, we isolated DNA with DirectPCR lysis Reagent (tail) (7Bioscience GmbH). PCR was performed with OneTaq® Quick-Load® 2X Master Mix (New England Biolabs) according to the manufacturer’s guidelines using the primers specified in Table S1.

All mouse strains were maintained on a C57BL/6 background for at least 6 generations. For experiments, randomly assigned 10 wk old males and females weighing at least 20 g were used. Mice were kept under pathogen-free barrier conditions.

### Cell culture, treatment and transfection

Human colorectal cancer cell lines RKO, LS513, LS174T (all harboring WTp53) and SW480 (harboring mutp53 R273H) were cultured in RPMI 1640 medium, isogenic HCT116 WTp53 and HCT115 p53null were cultured in McCoys medium, all supplemented with glutamine, 10% fetal bovine serum and penicillin/streptomycin and grown in a humidified atmosphere at 37°C with 5% CO_2_. All cell lines were regularly tested for mycoplasma contamination using the MycoAlert Mycoplasm detection kit (Lonza).

siRNAs were purchased from Ambion/Thermo Fisher Scientific (siRNAs are specified in Table 1) and transfected with Lipofectamine 2000 (Invitrogen). Nutlin-3a (BOC Biosciences), Palbociclib (Sigma), Idasanutlin (RG3788, SelleckChem), RG7112 (SelleckChem), RO-3306 (Sigma) and Roscovitine (Cell Signaling) were dissolved according to manufacturer’s guidelines and used as indicated.

**Table 1.**
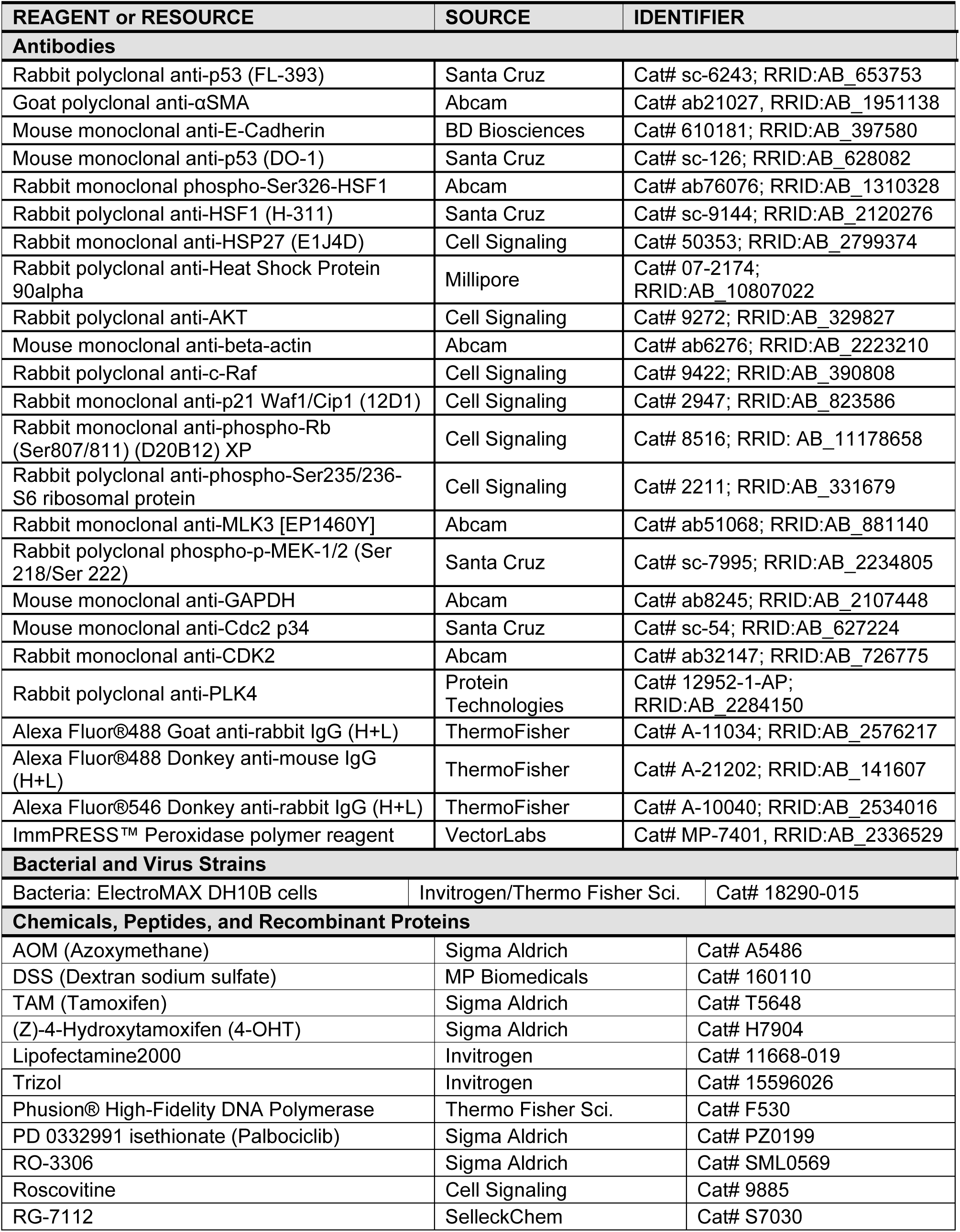
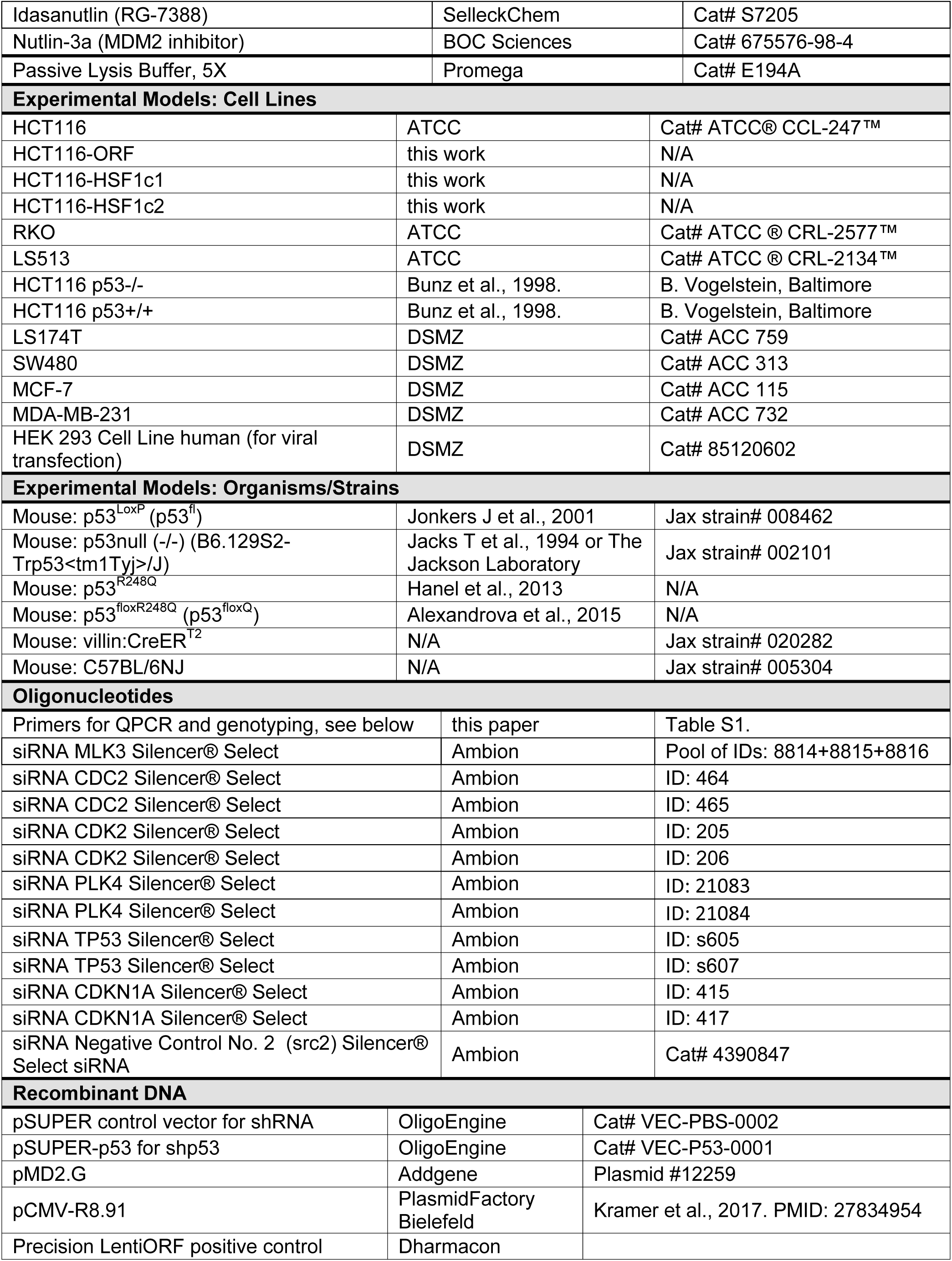
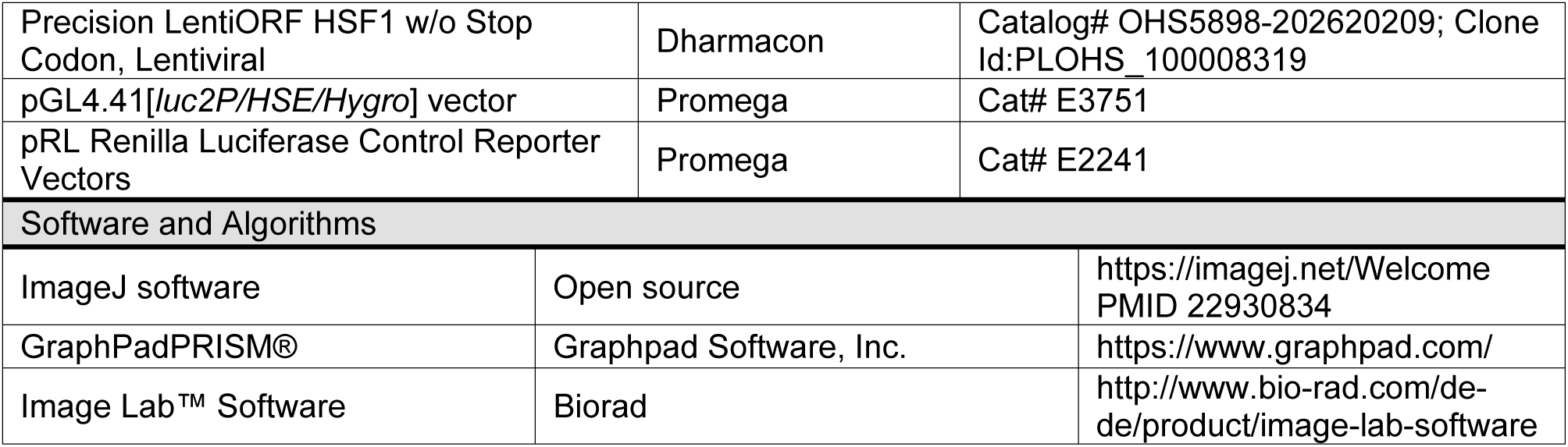
related to online Methods:: Reagents and Resources

For stable HSF1 expression in HCT116 cells, HEK-293 cells were co-transfected with lentiviral packaging vectors (*pMD2.G* from Addgene and *pCMV-R8.91* from PlasmidFactory Bielefeld) and the Precision LentiORF HSF1 lentiviral plasmid (Id:PLOHS_100008319) or a Precision control plasmid (Dharmacon). After standard lentivirus production, HCT116 cells were transduced in the presence of 8 µg/mL polybrene and cells were selected with Hygromycin for several days. Single cell clones were expanded and validated for HSF1 overexpression by immunostaining with phospho-Ser326 HSF1 (Abcam). Cell clones (HSF1c1, HSF1c2 and ORF control) were cultured in McCoys medium and supplemented as described above.

### CRC induction, colonoscopy and treatment

Murine colorectal carcinoma (CRC) was induced by a single intraperitoneal injection of the colon-selective carcinogen Azoxymethane (AOM, 10 mg/kg in 0.9% sodium chloride, Sigma) at the age of 10 wks. After one week rest, an acute colitis was induced with 1.5% (in p53-deficient mice) or 1.8% (in p53-proficient mice) dextran sodium sulfate (DSS, MP Biomedicals) for 6 days in the drinking water.

Visualization of tumor growth by mini endoscopy/colonoscopy (Karl Storz GmbH) started 6 wks after AOM induction. Tumor sizes were scored according to the Becker & Neurath score ^6^. Briefly, tumor sizes are calculated relative to the width (luminal circumference) of the colon and scored as sizes 1–5 (S1-S5) with the following specifications: S1 = just detectable, S2 = 1/8 of the lumen, S3 = 1/4 of the lumen, S4 = 1/2 of the lumen and S5 > 1/2 of the lumen. Notably, between 6-8 wks post AOM approximately 80% of mice had at least one S3 tumor and at least three S2 tumors.

As described in Schulz-Heddergott et al., 2018, for analysis of *TP53*^R248Q^ mice with either a constitutive p53 wildtype (+) or KO (-) allele, we chose an endpoint type of analysis, ending at 12 wks after AOM in p53-proficient mice (at least one WTp53 allele), or at 10 wks after AOM in all p53-deficient mice (deleted or mutated). This design prevented loss of mice due to colonic obstruction, anal prolapse, or lymphoma development in p53-deficient mice.

For analysis of the inducible p53LOH mouse model we used the *TP53*^R248Q^ allele combined with the conditional floxed WTp53 allele (p53^fl^) to create heterozygous p53^Q/fl^; vilCreER^T2^ tumors. We specifically induced p53LOH after a defined endoscopy-verified tumor burden was reached (at least one S3 tumor in addition to at least three S2 tumors). After tumor verification, Tamoxifen (TAM, Sigma) was given by 7 serial intraperitoneal injections (1 mg daily per injection in a 1:10 ethanol/oil mixture) to activate the inducible recombinase (*villinCreER*^*T2*^) and cause p53LOH. Tumor growth was continued to be visualized by colonoscopy over 2 - 8 wks after LOH induction by TAM.

At endpoints all mice were euthanized and the entire colon and rectum were harvested. Colons were longitudinally opened, cleaned and displayed. Tumor numbers were counted and tumor sizes measured with a caliper. Tumor biopsies were taken from all mice. To ensure complete sampling of the organ, each colon/rectum was ‘swiss rolled’, fixed in 4% paraformaldehyde/PBS and bisected. Both halves were placed face down side-by-side into a single cassette for histologic processing, paraffin embedding and subsequent tissue analysis.

Nutlin-3a (BOC Biosciences) treatment was given by oral gavage with 150 mg/kg per dose over 3 consecutive days. Mice were sacrificed and colorectal tumors harvested 8 wks after the last treatment.

### Histological analysis

Standardized immunohistochemical stainings were performed on murine formalin-fixed paraffin-embedded (FFPE) tissues. The following primary antibodies were used: p53 FL393 (Santa Cruz, sc-6243), pan-Cytokeratin (Abcam, ab9377) and α-smooth muscle actin/SMA (Abcam, ab21027). The ImmPRESS™ Peroxidase polymer reagent based on 3, 3-diaminobenzidine (DAB, Vectorlabs), or Alexa Fluor®488-coupled and Alexa Fluor®647-coupled secondary antibodies (immunofluorescence) were used as detection systems. Hematoxylin (DAB) or DAPI (immunofluorescence) were used as counterstains.

To define invasive mouse CRC tumor stages we used the following definition: cancer grown through the muscularis mucosae into the submucosa (=T1), cancer grown through the muscularis mucosae and submucosa into the muscularis propria (=T2), cancer grown into the outermost layers of the colon or rectum and reaching the serosa (=T3). No spread to nearby lymph nodes or distant metastasis were overserved.

### Immunoblots

Whole cell protein lysates were prepared with RIPA buffer (1% TritonX-100, 1% Desoxycholate, 0.1% SDS, 150mM NaCl, 10mM EDTA, 20mM Tris-HCl pH7.5 and complete protease inhibitor mix, Roche). Tumor tissues were minced and lysed with RIPA buffer followed by sonication. After centrifugation, protein concentrations were determined by BCA protein assay (Pierce). Equal amounts of protein lysates were separated by SDS-polyacrylamide gel electrophoresis (PAGE), transferred onto nitrocellulose membranes (Millipore), blocked with 5% milk and probed with the following antibodies: murine p53 (CM5, Vector Laboratories), human p53 (DO-1, Santa Cruz sc-126), total HSF1, pMEK1 and CDK1 (all Santa Cruz), HSP90α (Millipore), HSP27, AKT, cRAF, Bcl-Xl, CDKN1A/p21, phospho-RB and phospho-S6 (all Cell Signaling), MLK3, phospho-Ser326 HSF1 and CDK2 (all Abcam), PLK4 (Protein Technologies), GAPDH and beta-Actin (both Abcam). Detailed information of antibodies are listed in Table 1.

### Quantitative PCR

Total RNA from cells, tumor tissues or organoids was isolated using the Trizol reagent following manufacturers’ guideline (Invitrogen/Thermo Fisher Scientific). Tumor tissues were first homogenized using a homogenizer (T10 basic ULTRA-TURRAX). Equal amounts of RNA were reverse-transcribed (M-MuLV Reverse Transcriptase, NEB), and quantitative real-time PCR (qRT-PCR) analysis was performed using a qPCR Master-Mix (75 mM Tris-HCl pH 8.8, 20 mM (NH_4_)_2_SO_4_, 0.01% Tween-20, 3 mM MgCl_2_, SYBR Green 1:80,000, 0.2 mM dNTPs, 20 U/ml Taq-polymerase, 0.25% TritonX-100, 300mM Trehalose). Primers are specified in Table S1.

### Dual Luciferase Reporter (DLR) Assay

HSF1 firefly luciferase plasmids harboring seven HSE elements (pGL4.41[*luc2P*/HSE/Hygro] vector) and the pRL (Renilla) luciferase reporter plasmid (pRL-TK) were purchased from Promega. Cells were seeded and 24 hrs later were co-transfected with 100 ng HSF1*Luc* plasmids and 200 ng pRL-TK plasmid using Lipofectamine 2000 (Invitrogen). 48 hrs post-transfection, cells were treated with Nutlin as indicated and firefly luciferase and Renilla luciferase activities were measured using a Dual Luciferase Assay. Briefly, cells were lysed with PLB (Passive Lysis Buffer, 5X E194A) and incubated for 15 min. Supernatants were first incubated and measured with firefly luciferase buffer (25 mM Glycylglycine, 15 mM K2HPO4, 4 mM EGTA pH 8.0, 15 mM MgSO_4_, 4 mM ATP pH 7.0, 1.25 mM DTT, 0.1 mM CoA, 80 µM Luciferin) and then with Renilla luciferase buffer (1,1 M NaCl, 2.2 mM Na_2_EDTA, 0.22 M K2HPO_4_ pH 5.1, 0.5 mg/ml BSA, 1.5 mM NaN3, 1.5 µM Coelenterazine). Relative light units (RLUs) were measured in a Luminometer Berthold Centro LB 960 plate reader. Values were normalized to Renilla activity and relativized to the control treatment.

### Murine organoids, media, culturing and treatment

For preparation of organoid media, HEK293T cells stably expressing mRspondin or mNoggin (kindly provided by Dr. Tiago De Oliveira), or mWnt3a cells were cultured in DMEM (Gibco) supplemented with GlutaMAX™ (Gibco), 10% FBS (Merck), Penicillin-Streptomycin (10,000 U/mL, Gibco) and Sodium Pyruvate (Gibco) in a humidified atmosphere at 37°C with 5 % CO_2_. For HEK293T mRpondin-I 300 µg/mL Zeocin (InvivoGen, #ant-zn-05) and for HEK293T mNoggin 500 µg/mL G418 (Geneticin, InvivoGen, #ant-gn-1) were added to the medium during cultivation. After HEK293 cell expansions, culturing media were replaced by conditioned medium (CM) containing Advanced DMEM/F-12 (Gibco) supplemented with GlutaMAX™ (Gibco), Penicillin-Streptomycin (10,000 U/mL, Gibco) and 10mM HEPES (Gibco). 50 mL of CM were added per 175 cm^2^ flask and HEK293 cells allowed to grow for one week. Each CM media were sterilly filtered and aliquoted. Since mRspondin-I and mNoggin proteins are each fused to an Fc-tag, the quality of each batch was tested by Dot-blot analysis. Organoid media was composed of 50% CM Wnt3a, 20% CM mNoggin, 10% CM mRspondin-I, N2 and B27 (both Gibco), 5 µM CHIR 99021 (Axon Medchem), 3.4 µg/mL ROCK inhibitor (Y-27632), 500 nM A83-01, 10 mM Nicotinamide (Sigma-Aldrich), 80 µM N-Acetyl-L-Cysteine (all Sigma-Aldrich), and 200 ng/mL rmEGF (ImmunoTools).

For organoid preparation, tumor-harboring mice were sacrificed and the colons harvested. Tumors were dissected, washed and minced, and incubated with 2 mg/mL Collagenase type I solution (Gibco, dissolved in Advanced DMEM/F12) at 37°C for 30 min, while pipetting up and down every 10 min to dissociate the tumors. Small tumor fragments were transferred into a new Falcon tube using a cell strainer (100 µm mesh size). Fragments were centrifuged and washed with Advanced DMEM/F12. After centrifugation, tumor fragments were resuspended in cold Matrigel (Corning) and plated as gel drops on culture plates. After Matrigel polymerization at 37°C, organoids were cultured in organoid media and cultivated in a humidified atmosphere at 37°C with 5% CO2. Medium was exchanged every 2-3 days. Splitting of organoids was performed when organoids started to accumulate dead cells in the lumen (approx. once a week). To this end, organoids were recovered from Matrigel and disrupted manually by pipetting using 1 ml blue tips. For enzymatic dissociation, organoids were incubated with 0.25% trypsin at 37 °C for 10 min, washed with Advanced DMEM/F12, centrifuged and cultured as described above. Experiments with murine colonic organoids were done between passage 3 and 8. p53LOH was induced with 1 µM 4OHT (Sigma) for 24 - 48 hrs (as indicated in the figure legends) in CHIR 99021-free and Rock-free organoid media.

### Immunofluorescence staining of organoids

Organoids were fixed within Matrigel domes with 2% / 0.1% Paraformaldehyde/Glutaraldehyde /PBS for 30 min. After intensively washing steps with PBS, gel domes with fixed organoids were removed from the plate and transferred into a tube. Sucrose infiltration was started with 20% sucrose / PBS, followed by 40 % sucrose / PBS, each incubated over night or longer at 4°C until the domes settled down. After sucrose infiltration, organoids were embedded in TissueTEK (Tissue-Tek® O.C.T™ Compound) and 10 µM cryo-sections were cut. Sections were air-dried for 30 min at RT, pre-wetted with PBS and quenched with 10 mM NaBH4 / PBS twice for 5 minutes at room temperature each time. After washing steps, samples were permeabilised with 0.1% TritonX-100 / PBS for 10 min at RT and blocked with 10% FBS / 1% BSA / PBS for 1 hour. For staining, samples were co-incubated with the p53 antibody FL393 (Santa Cruz) and E-Cadherin (BD Biosciences) overnight at 4°C. Primary antibodies were detected by AlexaFluor488- and AlexaFluor647-conjugated secondary antibodies (Molecular Probes). Organoids were DAPI counterstained and mounted in Fluoromount media (DAKO). Images were taken using a standard fluorescence microscope (Carl Zeiss AG) with the ZEN imaging program from Zeiss. Figures were further prepared using Adobe Photoshop software.

### Analysis of human patient TCGA data

We used TCGA (The Cancer Genome Atlas) colorectal cancer (COARDREAD) and breast cancer (BRCA) databases in this analysis. Human genomic data including RNA expression, DNA copy number alteration, gene mutation, and clinical information was downloaded from cBioPortal for cancer genomics (http://www.cbioportal.org). Study names: Colorectal adenocarcinoma (TCGA, PanCancer Atlas, 594 total samples) and Breast Invasive Carcinoma (TCGA, PanCancer Atlas, 1084 total samples). TP53 wild type (WTp53) group are those samples without TP53 mutations. TP53 missense mutant group was samples with TP53 missense mutations (MS), and TP53 LOF group was determined by samples with all TP53 mutations (MS, missense; FS, frameshift; NS, nonsense). To identify tumors harboring p53LOH, we selected samples that had both a mutated TP53 gene and a shallow deletion in DNA copy number. The list of HSF1 target genes was chosen from Mendillo et al.^7^ We compared the expression values (by RNAseq) of HSF1 target genes from mutant p53/p53LOH tumors with samples that harbored wildtype TP53 (TP53^+/+^). Further we applied survival analysis to check patients with a missense TP53 mutation (MS p53) and a p53LOH compared to a WTp53 patient group. R language (The R Project for Statistical Computing, https://www.r-project.org) was used in the analysis. R package “gplots” was used to generate heatmaps. R package “survival” were used for survival analysis, including calculating log-rank p-values and generating Kaplan-Meier curves.

## QUANTIFICATION AND STATISTICAL ANALYSIS

Statistics of each experiment such as number of animals, number of tumors, biological replicates, technical replicates, precision measures (mean and ±SEM) and the statistical tests used for significance are provided in the figures and figure legends.

Unpaired Student’s t test was used to calculate the p values for comparisons of tumor numbers and sizes and mRNA expression levels.

Densitometric measurements for quantification of immunoblot bands were done with the gel analysis software Image Lab™ (BioRad) and normalized to loading controls.

The following designations for levels of significance were used within this manuscript: p* = 0.05; p** = 0.01; p*** = 0.001; ns, not significant.

---

**Table S2.**
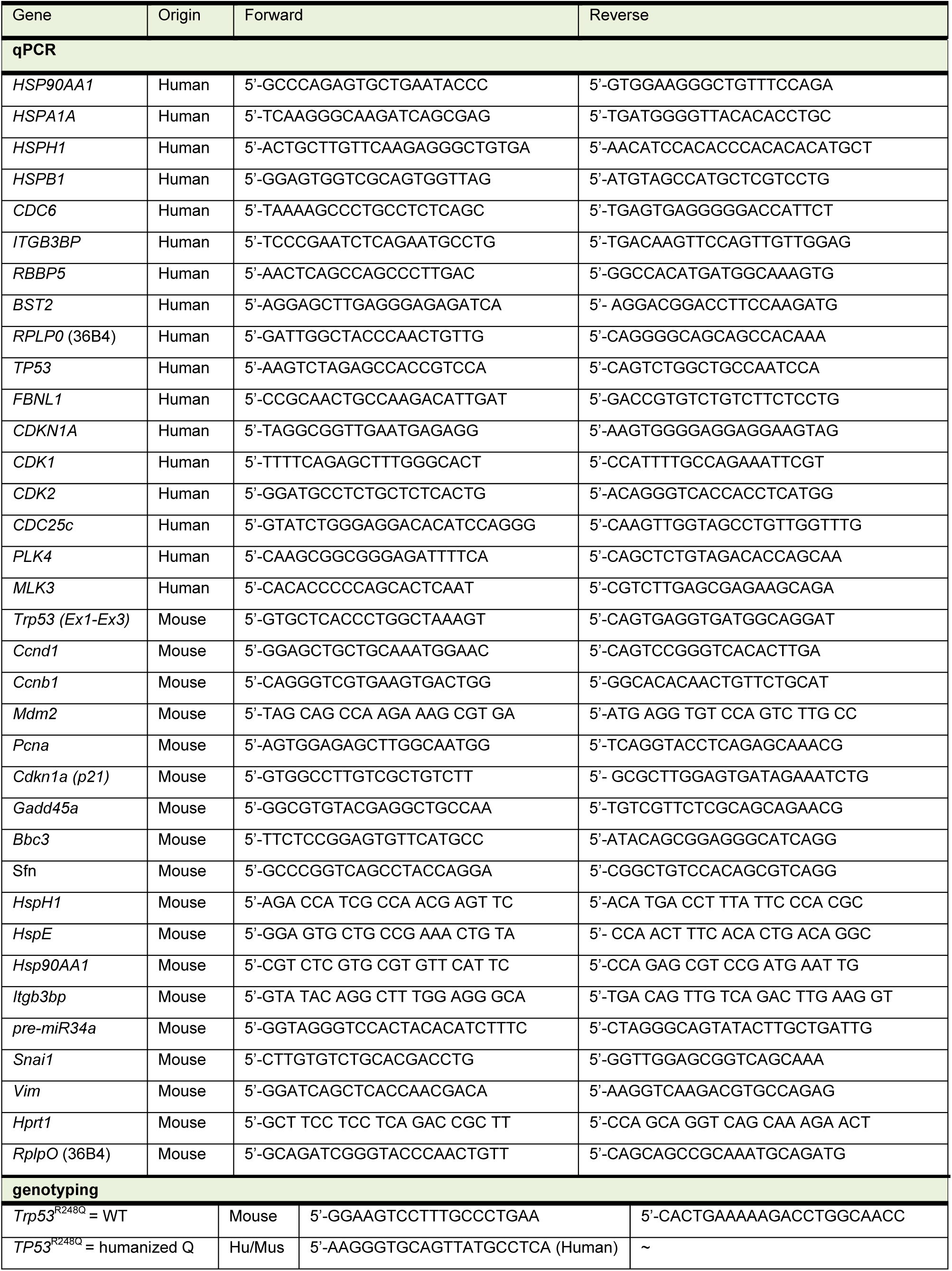
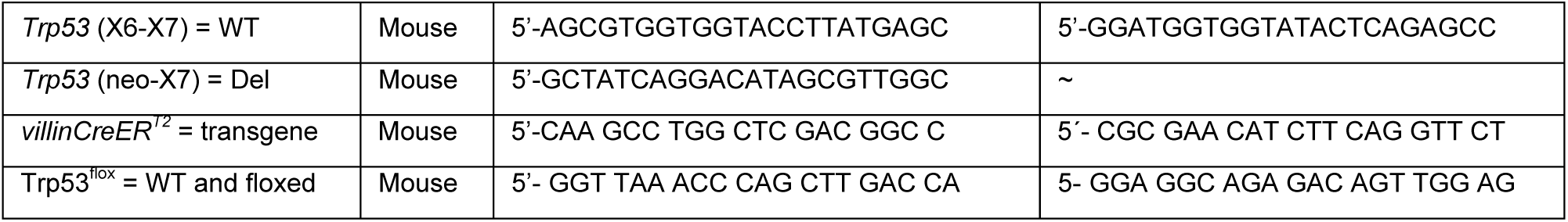
related to online Methods: Primers for qPCR and genotyping.

## Notes

### Competing Interest Statement

The authors have declared no competing interest.

## References

1. Fearon, E.R. & Vogelstein, B. A genetic model for colorectal tumorigenesis. Cell 61, 759–767 (1990).

2. Levine, A.J. & Oren, M. The first 30 years of p53: growing ever more complex. Nat Rev Cancer 9, 749–758 (2009).

3. Walerych, D., Lisek, K. & Del Sal, G. Mutant p53: One, No One, and One Hundred Thousand. Front Oncol 5, 289 (2015).

4. Brosh, R. & Rotter, V. When mutants gain new powers: news from the mutant p53 field. Nat Rev Cancer 9, 701–713 (2009).

5. Bykov, V.J.N., Eriksson, S.E., Bianchi, J. & Wiman, K.G. Targeting mutant p53 for efficient cancer therapy. Nat Rev Cancer 18, 89–102 (2018).

6. Nakayama, M. & Oshima, M. Mutant p53 in colon cancer. J Mol Cell Biol 11, 267–276 (2019).

7. Schwitalla, S. et al. Loss of p53 in enterocytes generates an inflammatory microenvironment enabling invasion and lymph node metastasis of carcinogen-induced colorectal tumors. Cancer Cell 23, 93–106 (2013).

8. Cooks, T. et al. Mutant p53 prolongs NF-kappaB activation and promotes chronic inflammation and inflammation-associated colorectal cancer. Cancer Cell 23, 634–646 (2013).

9. Schulz-Heddergott, R. et al. Therapeutic Ablation of Gain-of-Function Mutant p53 in Colorectal Cancer Inhibits Stat3-Mediated Tumor Growth and Invasion. Cancer Cell 34, 298–314 e297 (2018).

10. Cancer Genome Atlas, N. Comprehensive molecular characterization of human colon and rectal cancer. Nature 487, 330–337 (2012).

11. Goldstein, I. et al. Understanding wild-type and mutant p53 activities in human cancer: new landmarks on the way to targeted therapies. Cancer Gene Ther 18, 2–11 (2011).

12. Joerger, A.C. & Fersht, A.R. Structural biology of the tumor suppressor p53. Annu Rev Biochem 77, 557–582 (2008).

13. Olivier, M., Hollstein, M. & Hainaut, P. TP53 mutations in human cancers: origins, consequences, and clinical use. Cold Spring Harb Perspect Biol 2, a001008 (2010).

14. Schulz-Heddergott, R. & Moll, U.M. Gain-of-Function (GOF) Mutant p53 as Actionable Therapeutic Target. Cancers (Basel) 10 (2018).

15. Olive, K.P. et al. Mutant p53 gain of function in two mouse models of Li-Fraumeni syndrome. Cell 119, 847–860 (2004).

16. Terzian, T. et al. The inherent instability of mutant p53 is alleviated by Mdm2 or p16INK4a loss. Genes Dev 22, 1337–1344 (2008).

17. Hanel, W. et al. Two hot spot mutant p53 mouse models display differential gain of function in tumorigenesis. Cell Death Differ 20, 898–909 (2013).

18. Lang, G.A. et al. Gain of function of a p53 hot spot mutation in a mouse model of Li-Fraumeni syndrome. Cell 119, 861–872 (2004).

19. Nakayama, M. et al. Intestinal cancer progression by mutant p53 through the acquisition of invasiveness associated with complex glandular formation. Oncogene 36, 5885–5896 (2017).

20. Stein, Y., Rotter, V. & Aloni-Grinstein, R. Gain-of-Function Mutant p53: All the Roads Lead to Tumorigenesis. Int J Mol Sci 20 (2019).

21. Freed-Pastor, W.A. & Prives, C. Mutant p53: one name, many proteins. Genes Dev 26, 1268–1286 (2012).

22. Kim, M.P. & Lozano, G. Mutant p53 partners in crime. Cell Death Differ 25, 161–168 (2018).

23. Bellazzo, A., Sicari, D., Valentino, E., Del Sal, G. & Collavin, L. Complexes formed by mutant p53 and their roles in breast cancer. Breast Cancer (Dove Med Press) 10, 101–112 (2018).

24. Muller, P.A.J. & Vousden, K.H. Mutant p53 in Cancer: New Functions and Therapeutic Opportunities. Cancer Cell 25, 304–317 (2014).

25. Pfister, N.T. & Prives, C. Transcriptional Regulation by Wild-Type and Cancer-Related Mutant Forms of p53. Cold Spring Harb Perspect Med 7 (2017).

26. Zhang, Y. et al. Somatic Trp53 mutations differentially drive breast cancer and evolution of metastases. Nat Commun 9, 3953 (2018).

27. Blagosklonny, M.V., Toretsky, J., Bohen, S. & Neckers, L. Mutant conformation of p53 translated in vitro or in vivo requires functional HSP90. Proc Natl Acad Sci U S A 93, 8379–8383 (1996).

28. Whitesell, L., Sutphin, P.D., Pulcini, E.J., Martinez, J.D. & Cook, P.H. The physical association of multiple molecular chaperone proteins with mutant p53 is altered by geldanamycin, an hsp90-binding agent. Mol Cell Biol 18, 1517–1524 (1998).

29. Muller, P., Hrstka, R., Coomber, D., Lane, D.P. & Vojtesek, B. Chaperone-dependent stabilization and degradation of p53 mutants. Oncogene 27, 3371–3383 (2008).

30. Ingallina, E. et al. Mechanical cues control mutant p53 stability through a mevalonate-RhoA axis. Nat Cell Biol 20, 28–35 (2018).

31. Lee, M.K. et al. Cell-type, dose, and mutation-type specificity dictate mutant p53 functions in vivo. Cancer Cell 22, 751–764 (2012).

32. Li, D., Marchenko, N.D. & Moll, U.M. SAHA shows preferential cytotoxicity in mutant p53 cancer cells by destabilizing mutant p53 through inhibition of the HDAC6-Hsp90 chaperone axis. Cell Death Differ 18, 1904–1913 (2011).

33. Li, D. et al. Functional inactivation of endogenous MDM2 and CHIP by HSP90 causes aberrant stabilization of mutant p53 in human cancer cells. Mol Cancer Res 9, 577–588 (2011).

34. Anckar, J. & Sistonen, L. Regulation of HSF1 function in the heat stress response: implications in aging and disease. Annu Rev Biochem 80, 1089–1115 (2011).

35. Gomez-Pastor, R., Burchfiel, E.T. & Thiele, D.J. Regulation of heat shock transcription factors and their roles in physiology and disease. Nat Rev Mol Cell Biol 19, 4–19 (2018).

36. Whitesell, L. & Lindquist, S. Inhibiting the transcription factor HSF1 as an anticancer strategy. Expert Opin Ther Targets 13, 469–478 (2009).

37. Dai, C., Whitesell, L., Rogers, A.B. & Lindquist, S. Heat shock factor 1 is a powerful multifaceted modifier of carcinogenesis. Cell 130, 1005–1018 (2007).

38. Miyata, Y., Nakamoto, H. & Neckers, L. The therapeutic target Hsp90 and cancer hallmarks. Curr Pharm Des 19, 347–365 (2013).

39. Mendillo, M.L. et al. HSF1 drives a transcriptional program distinct from heat shock to support highly malignant human cancers. Cell 150, 549–562 (2012).

40. Toma-Jonik, A., Vydra, N., Janus, P. & Widlak, W. Interplay between HSF1 and p53 signaling pathways in cancer initiation and progression: non-oncogene and oncogene addiction. Cell Oncol (Dordr) 42, 579–589 (2019).

41. Alexandrova, E.M. et al. p53 loss-of-heterozygosity is a necessary prerequisite for mutant p53 stabilization and gain-of-function in vivo. Cell Death Dis 8, e2661 (2017).

42. Hingorani, S.R. et al. Trp53R172H and KrasG12D cooperate to promote chromosomal instability and widely metastatic pancreatic ductal adenocarcinoma in mice. Cancer Cell 7, 469–483 (2005).

43. Baker, S.J. et al. Chromosome 17 deletions and p53 gene mutations in colorectal carcinomas. Science 244, 217–221 (1989).

44. Parikh, N. et al. Effects of TP53 mutational status on gene expression patterns across 10 human cancer types. J Pathol 232, 522–533 (2014).

45. Jackson, E.L. et al. The differential effects of mutant p53 alleles on advanced murine lung cancer. Cancer Res 65, 10280–10288 (2005).

46. Donehower, L.A. et al. Integrated Analysis of TP53 Gene and Pathway Alterations in The Cancer Genome Atlas. Cell Rep 28, 1370–1384 e1375 (2019).

47. Donehower, L.A. et al. Integrated Analysis of TP53 Gene and Pathway Alterations in The Cancer Genome Atlas. Cell Rep 28, 3010 (2019).

48. Muzumdar, M.D. et al. Clonal dynamics following p53 loss of heterozygosity in Kras-driven cancers. Nat Commun 7, 12685 (2016).

49. Shetzer, Y. et al. The onset of p53 loss of heterozygosity is differentially induced in various stem cell types and may involve the loss of either allele. Cell Death Differ 21, 1419–1431 (2014).

50. Becker, C., Fantini, M.C. & Neurath, M.F. High resolution colonoscopy in live mice. Nat Protoc 1, 2900–2904 (2006).

51. Ghaleb, A., Yallowitz, A. & Marchenko, N. Irradiation induces p53 loss of heterozygosity in breast cancer expressing mutant p53. Commun Biol 2, 436 (2019).

52. Li, D., Yallowitz, A., Ozog, L. & Marchenko, N. A gain-of-function mutant p53-HSF1 feed forward circuit governs adaptation of cancer cells to proteotoxic stress. Cell Death Dis 5, e1194 (2014).

53. Esser, C., Scheffner, M. & Hohfeld, J. The chaperone-associated ubiquitin ligase CHIP is able to target p53 for proteasomal degradation. J Biol Chem 280, 27443–27448 (2005).

54. Iyer, S.V. et al. Allele-specific silencing of mutant p53 attenuates dominant-negative and gain-of-function activities. Oncotarget 7, 5401–5415 (2016).

55. Kern, S.E. et al. Oncogenic forms of p53 inhibit p53-regulated gene expression. Science 256, 827–830 (1992).

56. Sabapathy, K. The Contrived Mutant p53 Oncogene - Beyond Loss of Functions. Front Oncol 5, 276 (2015).

57. Shahbandi, A. & Jackson, J.G. Analysis across multiple tumor types provides no evidence that mutant p53 exerts dominant negative activity. NPJ Precis Oncol 3, 1 (2019).

58. Schulz, R. et al. HER2/ErbB2 activates HSF1 and thereby controls HSP90 clients including MIF in HER2-overexpressing breast cancer. Cell Death Dis 5, e980 (2014).

59. Polager, S. & Ginsberg, D. E2F - at the crossroads of life and death. Trends Cell Biol 18, 528–535 (2008).

60. Rattanasinchai, C. & Gallo, K.A. MLK3 Signaling in Cancer Invasion. Cancers (Basel) 8 (2016).

61. Hartkamp, J., Troppmair, J. & Rapp, U.R. The JNK/SAPK activator mixed lineage kinase 3 (MLK3) transforms NIH 3T3 cells in a MEK-dependent fashion. Cancer Res 59, 2195–2202 (1999).

62. Schroyer, A.L., Stimes, N.W., Abi Saab, W.F. & Chadee, D.N. MLK3 phosphorylation by ERK1/2 is required for oxidative stress-induced invasion of colorectal cancer cells. Oncogene 37, 1031–1040 (2018).

63. Tang, Z. et al. MEK guards proteome stability and inhibits tumor-suppressive amyloidogenesis via HSF1. Cell 160, 729–744 (2015).

64. Vydra, N. et al. 17beta-Estradiol Activates HSF1 via MAPK Signaling in ERalpha-Positive Breast Cancer Cells. Cancers (Basel) 11 (2019).

65. Hanahan, D. & Weinberg, R.A. Hallmarks of cancer: the next generation. Cell 144, 646–674 (2011).

66. Rokavec, M., Li, H., Jiang, L. & Hermeking, H. The p53/miR-34 axis in development and disease. J Mol Cell Biol 6, 214–230 (2014).

67. Kimura, A. et al. Nuclear heat shock protein 110 expression is associated with poor prognosis and chemotherapy resistance in gastric cancer. Oncotarget 7, 18415–18423 (2016).

68. Tanaka, T. Development of an inflammation-associated colorectal cancer model and its application for research on carcinogenesis and chemoprevention. Int J Inflam 2012, 658786 (2012).

69. Muller, P.A. et al. Mutant p53 drives invasion by promoting integrin recycling. Cell 139, 1327–1341 (2009).

70. Wawrzynow, B., Zylicz, A. & Zylicz, M. Chaperoning the guardian of the genome. The two-faced role of molecular chaperones in p53 tumor suppressor action. Biochim Biophys Acta 1869, 161–174 (2018).

71. King, F.W., Wawrzynow, A., Hohfeld, J. & Zylicz, M. Co-chaperones Bag-1, Hop and Hsp40 regulate Hsc70 and Hsp90 interactions with wild-type or mutant p53. EMBO J 20, 6297–6305 (2001).

72. Boettcher, S. et al. A dominant-negative effect drives selection of TP53 missense mutations in myeloid malignancies. Science 365, 599–604 (2019).

73. Lane, D.P. How to lose tumor suppression. Science 365, 539–540 (2019).

74. Sabapathy, K. & Lane, D.P. Therapeutic targeting of p53: all mutants are equal, but some mutants are more equal than others. Nat Rev Clin Oncol 15, 13–30 (2018).

75. Weissmueller, S. et al. Mutant p53 drives pancreatic cancer metastasis through cell-autonomous PDGF receptor beta signaling. Cell 157, 382–394 (2014).

76. Sakai, E. et al. Combined Mutation of Apc, Kras, and Tgfbr2 Effectively Drives Metastasis of Intestinal Cancer. Cancer Res 78, 1334–1346 (2018).

77. Bolt, A.B., Papanikolaou, A., Delker, D.A., Wang, Q.S. & Rosenberg, D.W. Azoxymethane induces KI-ras activation in the tumor resistant AKR/J mouse colon. Mol Carcinog 27, 210–218 (2000).

78. Takeda, H. et al. Transposon mutagenesis identifies genes and evolutionary forces driving gastrointestinal tract tumor progression. Nat Genet 47, 142–150 (2015).

79. Logan, I.R. et al. Heat shock factor-1 modulates p53 activity in the transcriptional response to DNA damage. Nucleic Acids Res 37, 2962–2973 (2009).

80. Nitta, M., Okamura, H., Aizawa, S. & Yamaizumi, M. Heat shock induces transient p53-dependent cell cycle arrest at G1/S. Oncogene 15, 561–568 (1997).

81. Jin, X., Moskophidis, D., Hu, Y., Phillips, A. & Mivechi, N.F. Heat shock factor 1 deficiency via its downstream target gene alphaB-crystallin (Hspb5) impairs p53 degradation. J Cell Biochem 107, 504–515 (2009).

82. Muller, L., Schaupp, A., Walerych, D., Wegele, H. & Buchner, J. Hsp90 regulates the activity of wild type p53 under physiological and elevated temperatures. J Biol Chem 279, 48846–48854 (2004).

83. Walerych, D. et al. Hsp90 chaperones wild-type p53 tumor suppressor protein. J Biol Chem 279, 48836–48845 (2004).

84. Walerych, D. et al. Hsp70 molecular chaperones are required to support p53 tumor suppressor activity under stress conditions. Oncogene 28, 4284–4294 (2009).

85. Jacks, T. et al. Tumor spectrum analysis in p53-mutant mice. Curr Biol 4, 1–7 (1994).

## References

1. Hanel, W. et al. Two hot spot mutant p53 mouse models display differential gain of function in tumorigenesis. Cell Death Differ 20, 898–909 (2013).

2. Alexandrova, E.M. et al. Improving survival by exploiting tumour dependence on stabilized mutant p53 for treatment. Nature 523, 352–356 (2015).

3. Schulz-Heddergott, R. et al. Therapeutic Ablation of Gain-of-Function Mutant p53 in Colorectal Cancer Inhibits Stat3-Mediated Tumor Growth and Invasion. Cancer Cell 34, 298–314 e297 (2018).

4. Jonkers, J. et al. Synergistic tumor suppressor activity of BRCA2 and p53 in a conditional mouse model for breast cancer. Nat Genet 29, 418–425 (2001).

5. Jacks, T. et al. Tumor spectrum analysis in p53-mutant mice. Curr Biol 4, 1–7 (1994).

6. Becker, C., Fantini, M.C. & Neurath, M.F. High resolution colonoscopy in live mice. Nature protocols 1, 2900–2904 (2006).

7. Mendillo, M.L. et al. HSF1 drives a transcriptional program distinct from heat shock to support highly malignant human cancers. Cell 150, 549–562 (2012).

